# Structure of the hepatitis C virus E1E2 glycoprotein complex

**DOI:** 10.1101/2021.12.16.472992

**Authors:** Alba Torrents de la Peña, Kwinten Sliepen, Lisa Eshun-Wilson, Maddy Newby, Joel D. Allen, Sylvie Koekkoek, Ian Zon, Ana Chumbe, Max Crispin, Janke Schinkel, Gabriel C. Lander, Rogier W. Sanders, Andrew B. Ward

## Abstract

Hepatitis C virus (HCV) infection is a leading cause of chronic liver disease, cirrhosis, and hepatocellular carcinoma in humans, and afflicts more than 58 million people worldwide. The HCV envelope E1 and E2 glycoproteins are essential for viral entry and infection, and comprise the primary antigenic target for neutralizing antibody responses. The molecular mechanisms of E1E2 assembly, as well as how the E1E2 heterodimer binds broadly neutralizing antibodies, remains elusive. We present the cryo-electron microscopy (cryoEM) structure of the membrane-extracted full-length E1E2 heterodimer in complex with broadly neutralizing antibodies (bNAbs) AR4A, AT12009 and IGH505 at ∼3.5 Å resolution. We resolve the long sought-after interface between the E1 and E2 ectodomains and reveal how it is stabilized by hydrophobic interactions and glycans. This structure deepens our understanding of the HCV fusion glycoprotein and delivers a blueprint for the rational design of novel vaccine immunogens and anti-viral drugs.

## Introduction

Despite substantial demand for an hepatitis C virus (HCV) prophylactic vaccine, vaccine development has been limited by the extensive sequence diversity of the virus, and the lack of structural information on the conformationally heterogeneous envelope glycoprotein E1E2 complex^1,2^. Previous studies suggest that eliciting broadly neutralizing antibodies (bNAbs), which target E1E2 during infection, correlate with viral clearance and protection in humans ^3-5^, while passively administered bNAbs protect against infection in animal models^6-8^. These observations provide a motivation for the development of an HCV vaccine aimed at inducing bNAbs ^1^.

HCV is an enveloped, single strand, positive-sense RNA virus from the Flaviviridae family. The RNA genome encodes a single polyprotein that is processed by host and viral proteases into three structural and seven non-structural proteins ^9^. The E1 and E2 envelope proteins associate to form a glycoprotein complex located on the outside of the virus that drives entry into hepatocytes^9^. The E2 subunit includes the receptor-binding domain and engages scavenger-receptor class B1 (SR-B1) and the tetraspanin CD81, while E1 is assumed to be the fusogenic subunit because it contains the putative fusion peptide (pFP)^10-13^. Since the E1E2 complex is the only viral protein on the surface of the virus, it is the sole target for bNAbs and thus an attractive candidate for structure-based immunogen design.

High-resolution structure determination of the full-length E1E2 heterodimer has been hindered by intrinsic flexibility, conformational heterogeneity, disulfide bond scrambling and abundant glycosylation^2,14-17^. The glycan shield not only protects E1E2 from immune recognition, but also facilitates assembly and viral infection^17-19^. Structural information is currently limited to truncated versions of recombinant E1 or E2, or small peptides^20-28^. Moreover, antigenic region 4 (AR4), a highly desirable target for vaccines because it includes the epitopes of several broad and potent HCV bNAbs such as AR4A and AT1618, has eluded structural characterization^5,29^. While membrane-associated E1E2 displays AR4 and binds bNAb AR4A efficiently, soluble HCV E1E2 glycoprotein complex usually does not, suggesting that AR4 represents a metastable domain^19,30-32^ (Sliepen et al. in prep). Using an optimized expression and purification scheme, we discovered that the co-expression and binding of AR4A stabilized the assembly of the full-length E1E2 heterodimer, producing a promising sample for structure determination. We subsequently determined the cryoEM structure of the E1E2 heterodimer in complex with the fragment antigen binding (Fab) of AR4A, and the Fabs of the bNAbs IGH505 and AT12009, providing a molecular description of three key neutralizing epitopes that paves the way for structure-based vaccine design.

### Purification and overall fold of E1E2

The full-length hepatitis C virus envelope glycoprotein complex E1E2 described here is derived from the genotype 1a strain AMS0232, which was obtained from an HCV-infected individual enrolled in the MOSAIC cohort ^33^. The AMS0232-based pseudovirus (HCVpp) was more resistant to neutralization by polyclonal HCV-positive sera than the reference strain H77, but sensitive to AR4A, making it suitable for pursuing a complex with this bNAb (Fig. 1a, Extended Data Fig. 1a) (Chumbe et al. in prep).

**Fig. 1.**
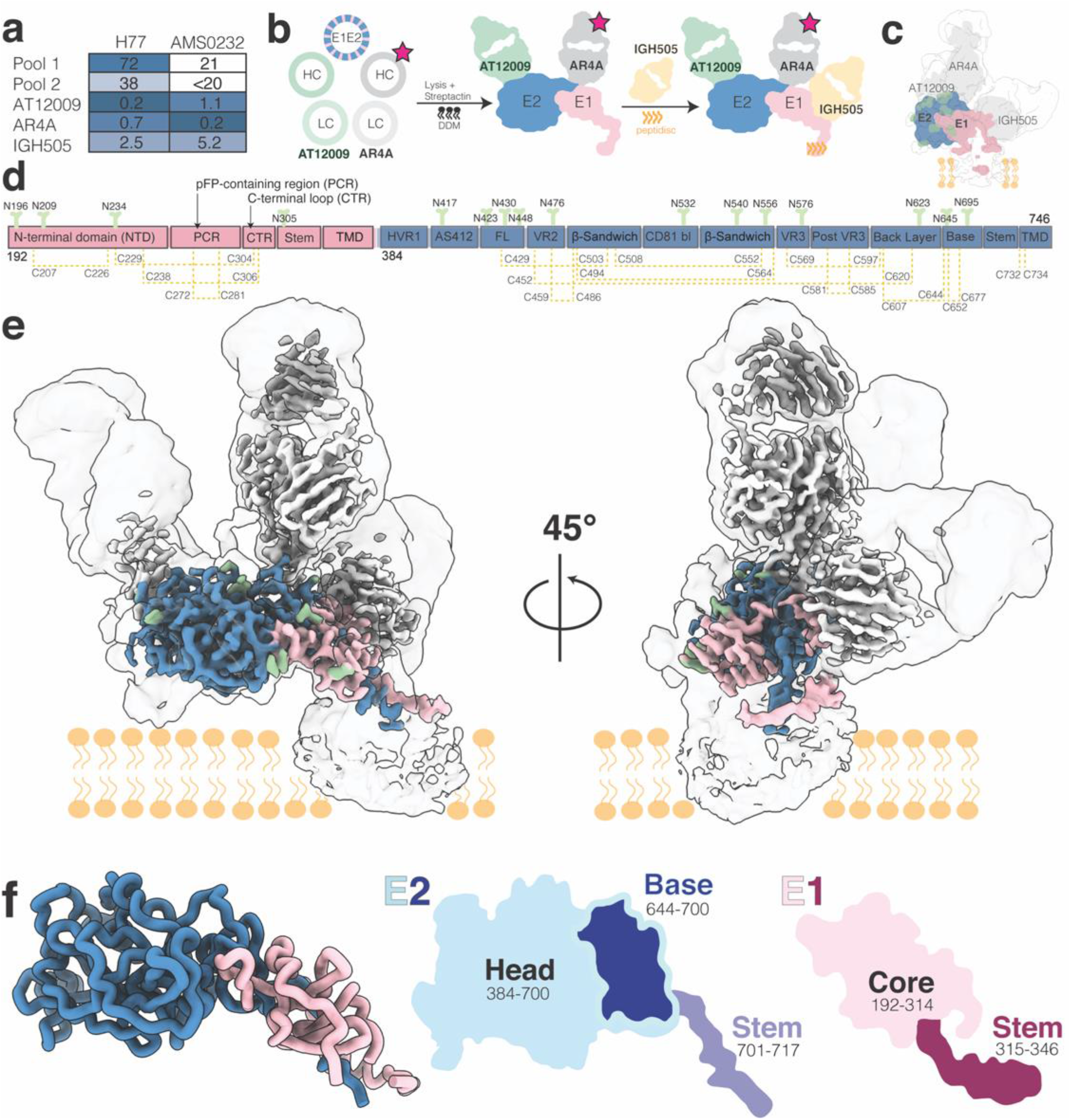
CryoEM structure of HCV envelope glycoprotein E1E2 heterodimer in complex with bNAbs AT12009, IGH505 and AR4A. (**a**) Sensitivity of AMS0232 pseudovirus to neutralization by polyclonal serum pools and bNAb AT12009, IGH505 and AR4A in comparison with H77 E1E2. The serum dilutions and antibody concentrations (in µ g/ml) at which HCV infectivity is inhibited by 50% (ID_50_ and IC_50_, respectively) are listed. Pseudovirus neutralization experiments were performed in duplicate or quadruplicate and values are the mean of two or three independent experiments. Darker shade indicates increased sensitivity. (**b**) Schematic representation of the purification of full-length HCV E1E2. Plasmids encoding full-length AMS0232 E1E2, strepII-tagged AR4A Fab (red star) and AT12009 crossFab were co-transfected in 293F cells and complexes were purified using Streptactin resin and embedded in peptidiscs, prior to the addition of IGH505 Fab. (**c**) Cartoon representation of the cryoEM map density of E1 and E2 in complex with AT12009, IGH505 and AR4A Fabs overlayed with the low-resolution cryoEM map at a threshold of 0.1 in ChimeraX. (**d**) Schematic representation of the full length E1E2 AMS0232 construct. The E1 subunit is shown in pink and the N-terminal domain (NTD), pFP containing region (pFP), C-terminal loop region (CTR), stem and transmembrane domain (TMD) regions are indicated. The E2 subunit is shown in blue with the different subdomains indicated: front layer, β-sandwich, CD81 binding site, back layer and base, hypervariable region 1 (HVR1), antigenic site 412 (AS412), variable region 2 (VR2) and variable region 3 (VR3), stem and TMD. N-linked glycans are shown in green and numbered by their respective Asn residues. Disulfide bonds are shown in yellow dashed lines and numbered accordingly. The color coding is preserved from (e) to (f). (**e**) CryoEM map showing the density of the full-length E1E2 in complex with the three bNAbs (AT12009, AR4A and IGH505). A rotated view by 45° is shown. (**f**) View of E1E2 heterodimer. A cartoon representation of the head and stem regions of E2 with the newly resolved base region highlighted as well as the core and stem regions of E1 are shown.

**Fig. 2.**
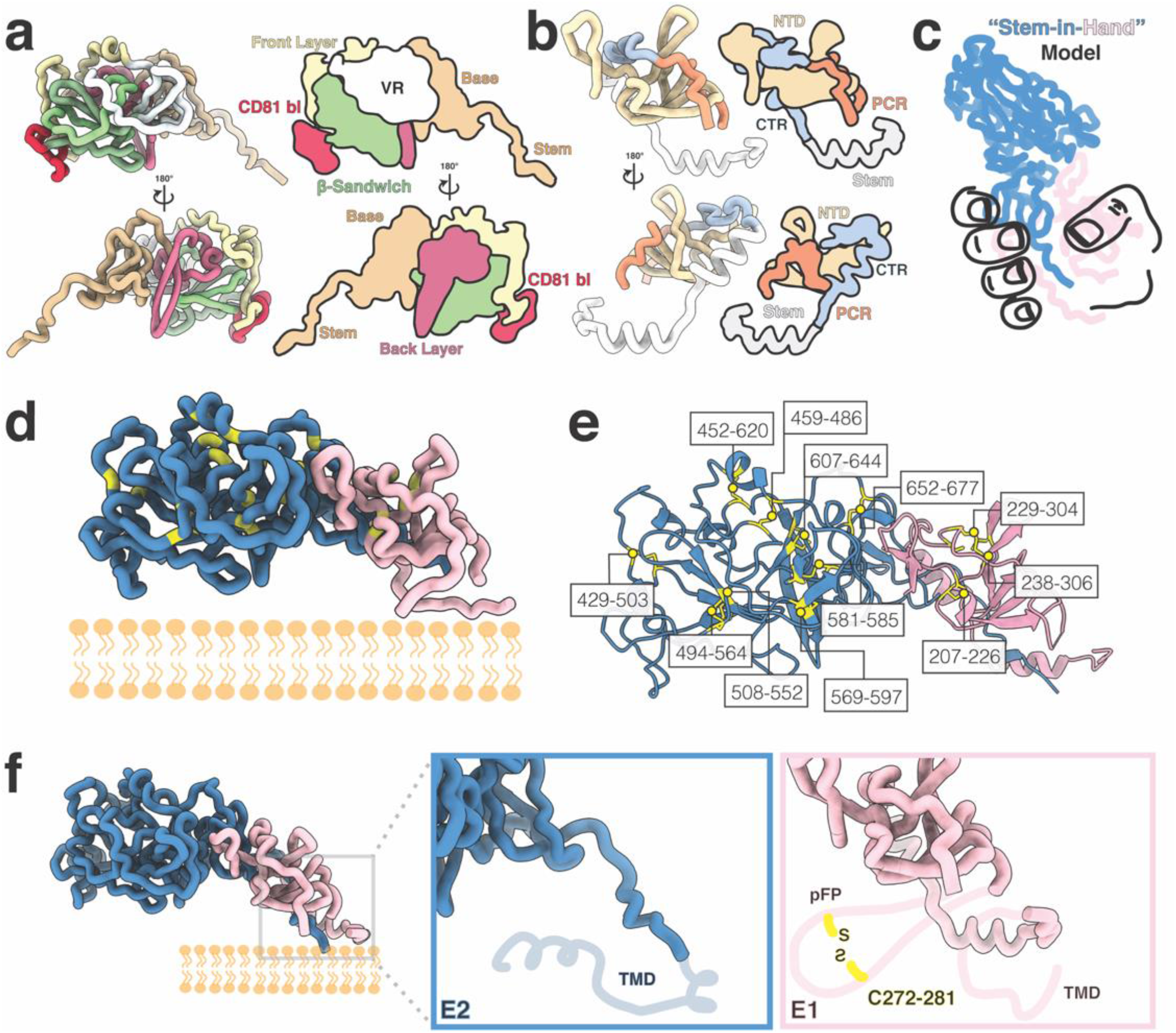
Subdomain organization and disulfide bonds of E1 and E2. View of E1E2 subdomains: (**a**) Each domain in E2 is colored and represented as licorice and cartoon. The E2 stem and base are shown in tan; followed by the back layer in magenta; β-sandwich in green; front layer in yellow; CD81 binding site in red; and all variable regions are shown in white. (**b**) The E1 N-terminal domain (NTD) is shown in yellow while pFP containing region (PCR) is shown in orange, the C-terminal loop region (CTR) in blue and the stem region is colored white. (**c**) A stem-in-hand model of E1 (hand) grasping the stem of E2. (**d**) The location of each cysteine in E1E2 is highlighted in yellow and further outlined and numbered in panel (**e**). (**f**) Close-ups of the E1E2 C-terminal region to highlight the missing regions in this highly flexible region: the transmembrane domain in E2 and two helices in E1 that comprise the E1 pFP-containing region and contain a conserved disulfide bond as well as a transmembrane domain (TMD) in E1 (indicated by the light pink cartoon line). The missing regions are depicted according to AlphaFold predictions.

We co-expressed strepII-tagged AR4A Fab with the E1E2 heterodimer and used StrepTactin purification to produce E1E2 in complex with AR4A (Fig. 1b). Binding of monoclonal Abs to E1E2 in complex with AR4A correlated with neutralization of the parental pseudovirus, suggesting that AR4A-extracted E1E2 is antigenically analogous to functional E1E2 (Extended Data Fig. 1b-d). For high-resolution cryo-EM studies, we co-expressed the full-length E1E2 glycoprotein complex with the AT12009 Fab^5^ and the strepII-tagged AR4A Fab to increase E1E2 stability. The complex was extracted from the membrane and reconstituted into peptidiscs, prior to the addition of the IGH505 Fab (Fig. 1b)^34^. To prevent mispairing of the AR4A and AT12009 heavy and light chains during synthesis, we used a CrossMAb version of the AT12009 Fab (CrossFab)^35^.

While we were able to isolate a biochemically well-behaved complex of E1E2 bound to Fabs, the relatively small size and flexibility of the complex presented substantial challenges for high resolution reconstruction. To overcome these challenges, we utilized 3D Variability Analysis in cryoSPARC^36^ to resolve both discrete and flexible conformations of the E1E2 heterodimer bound to three Fabs, resulting in a ∼3.5 Å resolution map (Extended Data Fig. 2). This structure was sufficient to model 51% of E1 and 82% of E2, including the interface of the two envelope glycoproteins, the epitopes of bNAbs AR4A, AT12009 and IGH505 and the glycan shield (Fig. 1c-e, Extended Data Fig. 3). We also resolved the majority of E1E2 complex, excluding the disordered hypervariable region 1 (HVR1) in E2 (384-419), antigenic site 412 (AS412, amino acids 411-419) in E2, the transmembrane domains (TMDs) in E1 and E2 (amino acids 349-382 and 718-746, respectively) and, lastly, the putative fusion peptide (pFP)-containing region (PCR) in E1 (residues 257-294) (Extended data Fig. 3, 4a). Although the E1 and E2 TMDs were unresolved, we combined the AlphaFold-predicted structure of these domains with our experimentally derived model to derive insights into the positioning of E1E2 relative to the membrane (Extended Data Fig. 3c-e).

**Fig. 3.**
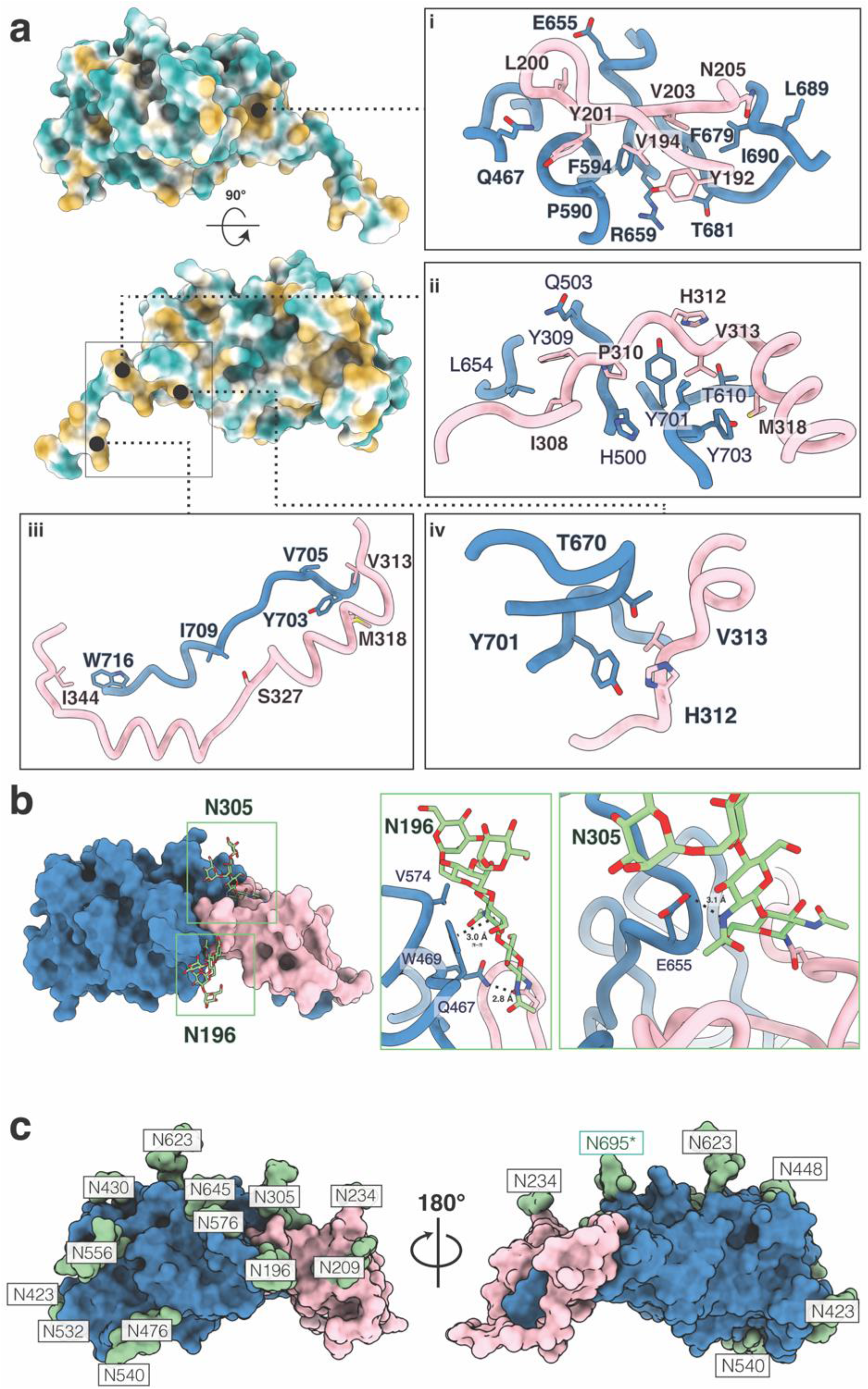
The E1E2 interface and glycan shield. (**a**) The novel E1E2 interface is stabilized by hydrophobic interactions. E2 is colored by hydrophobicity with green representing hydrophilic regions and yellow signifying hydrophobic patches. The first panel showcases a deep hydrophobic cavity in the base of E2 against which E1 packs. The following panels highlight additional hydrophobic interactions that we assert further stabilize the E1E2 interface. (**b**) Glycans buttress the E1E2 interface. Key interactions between glycans N196 and N305 are showcased. Glycan N196 is involved in hydrophobic interactions including a π-π stacking interaction with W469, as well as a salt bridge with Q467. N305 forms a stabilizing salt bridge with E655. (**c**) Surface representation of the E1 and E2 model showing the glycan sites in green with their respective Asn residues. The predominant glycoform identified by LC-MS at each PNGS was modelled using Coot^56^. An asterisk indicates that the glycan at position N695 uses a non-canonical N-glycosylation motif, NXV.

**Fig. 4.**
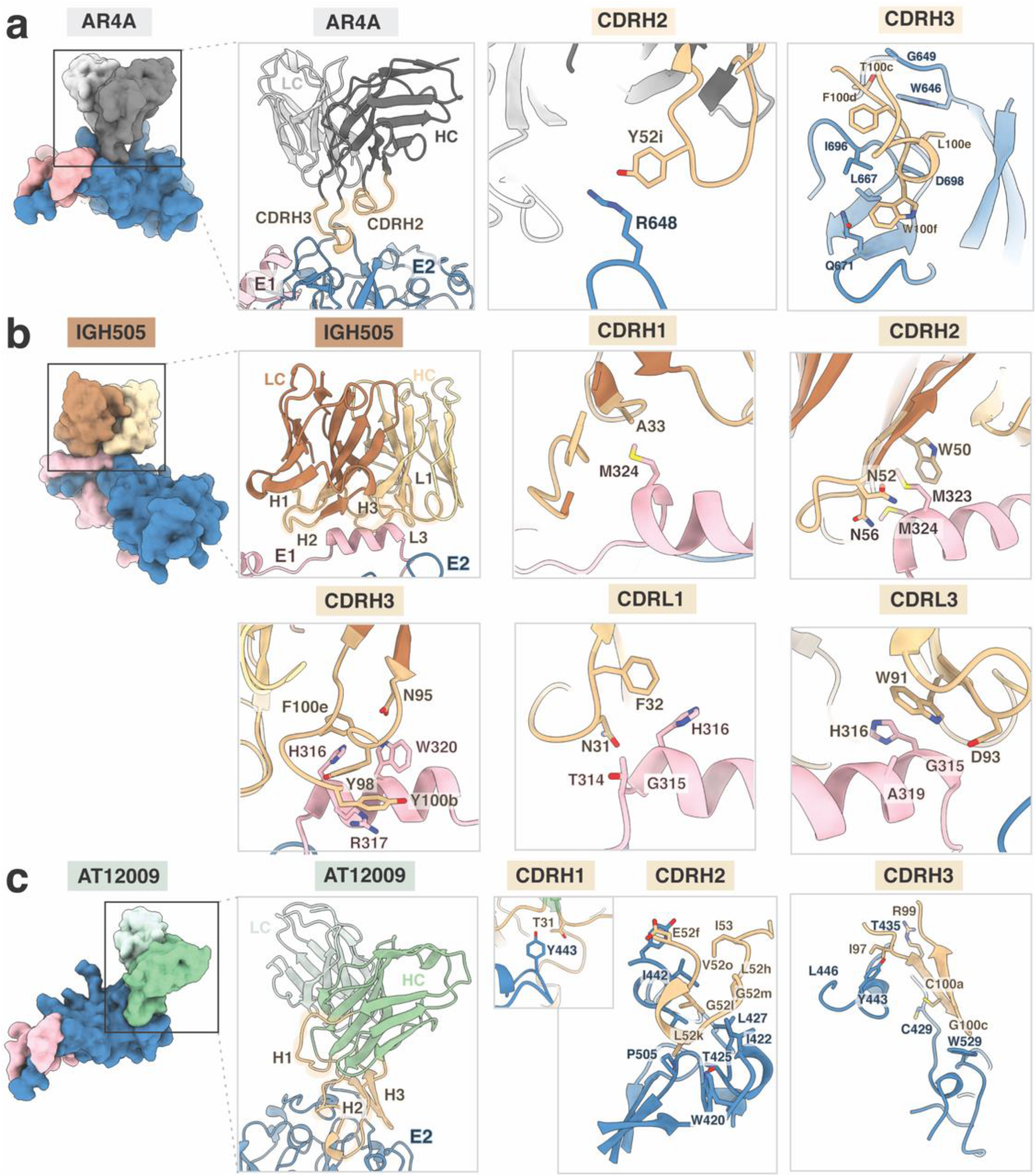
Structural definition of AR4A, IGH505 and AT12009 epitopes. (**a**) AR4A Fab recognizes protein elements in E2 (blue) near the interface with the E1 subunit (pink). Heavy and light chain are shown in dark and light gray, respectively and CDRH2 and CDRH3 are highlighted in tan. While the CDRH2 only interacts with the stem region of E2, the CDRH3 loop targets both the back layer and stem of E2. (**b**) IGH505 Fab interacts with the surface exposed α-helix in E1 (pink). Heavy and light chains are shown in brown and yellow, respectively and CDRH1-3 CDRL1 and 3 are highlighted in tan. IGH505 encases the conserved alpha helix in E1 (amino acids 310-328) using CDRH1-3 and CDRL1 and CDRL3 regions of the Fab. (**c**) AT12009 Fab binds the front layer of E2 (blue). Heavy and light chain are represented in green and light green, respectively and CDRH1-3 are highlighted in tan. All the CDRH loops interact with the front layer of E2, near the CD81 binding site. Epitopes on the E1 and E2 were defined as residues containing an atom within 4Å of the bound Fab and the amino acids present in the epitope are shown as sticks representations.

### Subdomain organization of E1 and E2

E2 contains three subdomains, termed the head, stem, and TMD. Our structure of the E2 head domain in the E1E2 complex agrees well with previously reported crystal structures of recombinant E2, including the most complete crystal structure (RMSD of 0.805 Å, PDBID: 6MEI) (Extended Data Fig. 4a). However, our E1E2 structure reveals two previously unresolved regions of E2: (1) an extended loop interspersed by a small anti-parallel β-sheet in the E2 head that we term the “base” (residues 645-700) and (2) the stem (residues 701-717) that connects the base with the TMD (amino acids 718 to 746; Fig. 1f, 2a). The E2 head consists of a central β-sandwich core (residues 484-517 and 535-568), a front layer and a back layer (residues 420-458 and 596-643, respectively), the apical CD81 binding site (amino acids 518-534), HVR1 (residues 384-410), AS412 (residues 411-419), variable regions 2 and 3 (VR2; residues 459-483, VR3; residues 569-579), and the newly resolved base (645-700).

Our structure also contains well-resolved density for the following regions in E1: the core (residues 192-314) and the stem (315-346) (Fig. 1f). The E1 core includes the N-terminal domain (NTD; 192-248), the pFP-containing region (PCR; 249-299) and a C-terminal loop region (CTR; 300-314) that connects the PCR to the stem (Fig. 1f, 2b). Intriguingly, the conformation of the E1 NTD in our cryo-EM structure differs substantially from that of a prior crystal structure of the soluble E1 NTD^26^ (PDB 4UOI), demonstrating that the presence of E2 is required for proper E1 folding. However, a prior crystal structure of 10 residues within the E1 stem (314-324) aligns well with our atomic model (310-328) (RMSD = 0.254 Å) (Extended Data Fig. 4a)^28^. The cryoEM structure also reveals the stem that connects the E1 core to its TMD anchoring E1 to the membrane (Fig. 1f, 2b; Extended Data Fig. 4a, c).

### Disulfide bond networks of E1 and E2

The cysteine residues in E1E2 are highly conserved, but disulfide bond patterns differ between recombinant E2 structures^20-22,37^, while the disulfide bond network in E1 has remained largely elusive^38^. Our cryoEM structure revealed that E2 is stabilized by nine disulfide bonds: C429-C503, C452-C620, C459-C486, C494-C564, C508-C552, C569-C597, C581-C585, C607-C644 and C652-C677 (Fig. 2d-f). Meanwhile, E1 is held together by four disulfide bonds: C207-C226, C229-C304, C238-C306 and C272-C281 (Fig. 2d-f). The E2 disulfide network is consistent with that of previous crystal structures of the E2 ectodomain, although the C652-C677 cysteine pair in the E2 base was missing from these structures^22,39^. In contrast, most recombinant E2 structures and a structure of E2 in complex with AR3C (PDB: 4MWF), the cysteines at positions C569, C581, C585 and C597 are paired differently compared to those in our cryoEM structure and other E2 crystal structures (Extended Data Fig. 3). We directly observed three disulfide bonds in E1 (C207-C226, C229-C304 and C238-C306) and AlphaFold predicted the presence of a fourth disulfide bridge between C272-C281, which resides between two amphipathic helices of the pFP. Given the proximity of the cysteine pairs C494-C564 to C508-C552, C452-C620 to C459-C486, and C581-C585 to C569-C597 and C607-C644 in E2, as well as the close proximity of the three observed disulfide bonds in E1, it is easy to understand how disulfide scrambling could occur, resulting in highly heterogeneous E1E2 proteins (Fig. 2e-f)^20,21,37,40^.

### The E1E2 interface

To illustrate the interface between E1 and E2, we posit a “stem-in-hand” model: the E1 wraps around the E2 stem and interacts with the base of E2 (Fig. 2c). The interactions between E1 and E2 are non-covalent as disulfide bonds are absent from the interface, which is consistent with earlier studies on intracellular E1E2^41,42^. Instead, the interface is lined with residues that mediate hydrophobic interactions and/or form hydrogen bonds. We observe a hydrophobic patch that lines the E2 stem to stabilize the E1E2 heterodimer interface (Fig. 3) The hydrophobic cavity on the E2 base involves residues F586, P590, F679, T681, L689, and I690 that interact with Y192, V194, Y201, and V203 on E1, while E655 and R659 form hydrogen bonds with L200 and G199 (Fig. 3a i). Within the hydrophobic stretch in E2 stalk, a region consisting of H500, L654, Y701, and Y703 interacts with I308, Y309, P310, H312, V313, M318 in E1 (Fig. 3a ii, iv) and the stretch of residues Y703, V705, I709 and W716 pair with E1 residues V313, M318, S327, and I344 (Fig. 3a iii).

The E1E2 interaction is further buttressed by two highly conserved glycans at N196 and N305 in E1 (Fig. 3b). The N305 glycan forms a salt bridge with E655 in E2, while the glycan at N196 forms multiple interactions, including a salt bridge with Q467, π–π stacking interactions with the aromatic ring of W469, and hydrophobic packing interactions with V574 (Fig. 3b). Both glycans are highly conserved (>99% among 505 HCV sequences), and their elimination reduced E1E2 complex formation and HCV pseudovirus infectivity, confirming their biological importance^18^.

### The E1E2 glycan shield

All potential *N*-glycosylation sites (PNGS) are located on one face of the E1E2 glycoprotein complex, while the opposite face, which is hydrophobic and highly conserved, does not contain PNGS (Extended Data Fig. 5a). These observations are consistent with an exposed neutralizing face subject to immune pressure, while the non-neutralizing face is likely inaccessible on the surface of the virion^23,24,43^. We detect density for glycans at all PNGS, except at N325 in E1, which is not glycosylated because of a proline at the fourth position within the N325 sequon^44^ (Extended Data Fig. 5c). Interestingly, while *N*-glycans are usually located at NXT/NXS sequons, we also discovered an *N*-glycan attached to a non-canonical NXV motif at N695 in E2 (Fig. 3c; Extended Data Fig. 5b). Site-specific glycan analysis of the E1E2 heterodimer in complex with AR4A and AT12009 sample using LC-MS confirmed the presence of glycans at all canonical PNGS except N325, as well as at the non-canonical N695 site, which was glycan-occupied in 25% of the sample (Fig. 3c, Extended Data Fig. 5b). Moreover, the E1E2 heterodimer was highly enriched in oligomannose-type glycans (Man_5_-_9_GlcNAc_2_), except for N234 (Extended Data Fig. 5b). The high oligomannose is likely due to the fact that the transmembrane domain of E1 is a signal for static retention in the ER as well as the purification from intracellular compartments, including the ER^45,46^.

### Definition of three bNAb epitopes

Importantly, our full-length E1E2 structure delivers a structural description of three bNAb epitopes in their full quaternary context, including the previously ill-defined epitopes in antigenic region 4 (AR4). Previous studies mapped the AR4A epitope using alanine-scanning mutagenesis, which suggested that AR4A recognizes an epitope on both E1 and E2^29,47^. However, our model shows that AR4A targets the back layer and base regions of E2 but does not engage E1 (Fig. 4a). AR4A contacts E2 almost exclusively using its 25 amino acid long CDRH3 via five hydrophobic interactions and four hydrogen bonds (AR4A - E2: T100c - G649, F100d - I696, L100e - D698, W646, P676; W100f - L667, Q671, I696) (Fig. 4a, Extended Data Fig. 6a, Extended Data Fig. 8). AR4A contains a nine amino acid insert in the CDRH2 (Extended Data Figure 6) and this allows Y52i to electrostatically interact with R648 in the back layer of E2 (Fig. 4). Notably, one glycan in E2 (N623) interacts with the N-terminal domain of the light chain of the Fab respectively (Extended Data Fig. 7a). The observation that AR4A does not bind directly to E1, but that mutations in E1 affect AR4A binding, suggest that the epitope of AR4A is metastable and requires E1 for stable presentation^29,47^.

bNAb IGH505 targets the surface-exposed conserved α-helix in E1, which is positioned against the CDRH3 loop and inserted between CDRH1, CDRH2, CDRL1, and CDRL3 loops with five CDR loops making contact with the epitope^48^. IGH505 targets H316 and W320 in E1, which bind to CDRH3, CDRL1, and CDRL3, likely through hydrogen bonds to D95 and π–π interactions with Y98, F100e, F32 and W91 (Fig. 4b, Extended Data Fig. 8). Also, M323 and M324, located at the C-terminal domain of the α-helix in E1, are within hydrogen bonding distance of A33 in CDRH1 and W50, and K58 in CDRH2 (Fig. 4b, Extended Data Fig. 8). While these latter interactions are not seen in the crystal structure of the E1 helix in complex with IGH526, both antibodies share similar footprints, indicating that they target an overlapping epitope at a very similar angle (Extended Data Fig. 6b and 7b)^28^.

The AT12009 bNAb targets the antigenic region 3 (AR3), which involves the front layer and the CD81 binding loop in E2. AT12009 contains the longest CDRH3 loop among all known AR3-targeting bNAbs (25 amino acids) and shares a similar footprint with the previously described antibodies HEPC3, HEPC74, AR3C, AR3A, HC11 and AR3X (Fig. 4c and Extended Data Fig. 7c). The sequence signature of these VH1-69-derived bNAbs is a CDRH3 that contains a β-hairpin-like structure stabilized by a disulfide bond (CxGGxC motif) (Extended Data Fig. 7d). The β-hairpin conformation adopted by the CDRH3 of AT12009 involves four hydrogen bond interactions between its CDRH3 and the front layer of E2 and CD81 binding loop: I97-A248, R99-T435, C100a-C429 and G100c-W529 (Fig 4c, Extended Data Fig. 7d, 8).

While the CDRH3 dominates the paratope of most of the AR3-targeting bNAbs isolated to date, the unusually long CDRH2 of 32 residues (Kabat numbering) contributes significantly to the interaction of AT12009 with the front layer of E2, with nine residues within hydrogen bond distance: E52f, G52l, G52m, L52h and I53; and burying similar surface area (493 Å^2^) than the CDRH3 (526 Å^2^) (Fig. 4c; Supplementary Table 2; Extended Data Fig. 7e, 8). A similarly ultralong CDRH2 (31 residues) is present in AR3X^22^. Collectively, these data give us comprehensive insights into the neutralizing face of E1 and E2 and facilitate structure-based vaccine design (extended Data Fig. 9).

## Discussion

The structure of the HCV envelope glycoprotein complex E1E2 proved challenging to resolve due to the need to co-express E1 and E2 and the propensity to form heterogeneous soluble aggregates even when expressed without the transmembrane domain^30,49-51^. Previous studies found that HCV virions mostly carry disulfide-linked E1E2 oligomers, leading to the assertion that the E1E2 interface was stabilized by disulfide bridges^45^. Other studies suggest that properly folded E1E2 heterodimers are noncovalently linked^42^. We demonstrate that the E1E2 interface is, in fact, stabilized by glycans and hydrophobic patches, and not by covalent disulfide bridges— providing insight into how to engineer stable immunogens for structure-based vaccine design^52-54^.

Co-expression of bNAb AR4A was key to stabilizing the metastable E1E2 complex, which we surmise is arrested in the pre-fusion conformation. AR4A efficiently neutralizes HCV *in vitro* and protects against infection *in vivo*. Its epitope proximal to the E1E2 interface likely stabilizes E1E2, therefore also likely blocking conformational changes associated with HCV entry^7,8,29^.

Our cryoEM structure elucidates the HCV E1E2 glycan shield that includes a rare glycan addition to a non-canonical NXV sequon, which was subsequently confirmed by mass spectrometry. Unlike other glycoproteins, where glycans are additive to the stability between interfaces, two glycans are necessary in stabilizing the E1E2 interface. The remaining glycans are distributed across one side of the E1E2 glycoprotein complex. Since antibodies bind to this side of E1E2, we refer to this as the neutralizing face of the glycoprotein complex. The opposite face of E1E2 lacks glycans and is primarily hydrophobic, and therefore may be involved in oligomerization and/or interaction with the viral lipid bilayer^55^. Overall, our cryoEM study provides a full molecular description of E1E2, including three bNAb epitopes, and provides a long sought-after blueprint for the design of a new generation of HCV glycoprotein immunogens and anti-viral drugs.

## Acknowledgements

We thank B. Anderson, H. L. Turner and M. Wu for cryoEM data collection support; C. Bowman and J.C. Ducom for computational support; A. Antanasijevic, G. Ozorowski, P. Sauer, D. Montiel-Garcia, E. Watson, B. Basanta, and J. Yang for supportive discussions. We thank Jelle Koopsen for the sequence alignment used for generating Extended Data Fig. 5a and Marit J. van Gils for supervising HCVpp assays. We also acknowledge The Scripps Research Institute CryoEM Facility and additional scientific resources at The Scripps Research Institute. We thank Steven Foung for donating the 212.10 and 212.15 antibodies and Tim Beaumont and Sabrina Merat for donating the AT12009 and AT1618 antibodies and antibody plasmids. R.W.S is a recipient of a Vici grant from the Netherlands Organization for Scientific Research (NWO). J.S is a recipient of a Vidi and Aspasia grant from the Netherlands Organization for Scientific Research (NWO, grant numbers 91719372 and 015.015.042). We thank the National Science Foundation Grant 2109312 to L.E.W., the Netherlands Organization for Scientific Research (NWO) Rubicon Grant 45219118 to A.T.d.l.P and the Amsterdam institute for Infection and Immunity Postdoctoral grant to K.S. Mass spectrometry (M.C.) was supported by Bill and Melinda Gates Foundation grant INV-008352/OPP1153692.

## Author Contributions

A.T.d.l.P. optimized sample preparation for cryoEM analysis and collected cryoEM data. L.E.-W assisted in cryoEM data collection and processed the data. A.T.d.l.P and L.E.-W built and refined the atomic models into the cryoEM maps. K.S. conceived the protein purification strategy and designed the constructs.. K.S., S.K., I.Z., A.C., performed protein purifications, ELISA experiments and neutralization assays. J.D.A. and M.N. ran site-specific glycosylation analysis. J.S. provided crucial materials. A.B.W., G.C.L., K.S. and R.W.S. provided guidance. A.T.d.l.P, L.E.-W., K.S., G.C.L., R.W.S and A.B.W wrote the paper. All authors contributed to the manuscript text by assisting in writing or providing feedback. R.W.S. and A.B.W. supervised the research.

## METHODS

### E1E2 and antibody constructs

The AMS0232 E1E2 sequence was obtained from an HIV-1 infected individual from the MOSAIC cohort study^33^. The full-length amino acid E1E2 sequence (amino acids 192-746, H77 numbering)^57^ was codon-optimized for expression in human cell lines and cloned into the pPPI4 expression vector downstream of a custom signal peptide (MDAMKRGLCCVLLLCGAVFVSVTG).

Amino acid sequences of the antibody variable heavy (VH) and light chains (VL) were codon-optimized and cloned into IgG1 in mammalian expression vectors. The StrepII-tagged heavy chain (HC) Fab of AR4A was generated by introducing a Gly-Gly-Ser-linker followed by a StrepII-tag directly downstream of the cysteine that pairs with the light chain (LC). For the production of the CrossFab, we generated an AT12009 VL-CH construct by fusing the VL to the heavy constant domain (CH) of IgG1 Fab and an AT12009 VH-CL construct by fusing the VH to the constant light (CL) domain separated by elbow regions as described by Schaefer and colleagues^35^. CDR loops and antibody amino acids are defined according to Kabat nomenclature and numbering scheme.

### Antibody and Fab production

All proteins were transiently transfected in suspension HEK293F cells (Invitrogen, cat no. R79009) maintained in Freestyle medium (Life Technologies). For antibody production, HEK293F were grown to a density of ∼1.0 × 10^6^ cells/ml and were co-transfected with heavy chain (HC) and (LC) plasmids (1:1) mixed with PEIMAX (Polysciences). After five days, the supernatant was harvested, filtered and the antibodies were purified using protein G or protein A agarose resin (Pierce) according to manufacturer’s instruction. The eluted antibodies were concentrated in PBS and injected in a Superdex200 Increase column equilibrated in PBS. Peak fractions were collected and concentrated with Vivaspin6 10-kDa cutoff filters. To produce IGH505 Fab, we incubated purified IGH505 IgG1 with papain-resin (Thermo Fisher) in digestion buffer (PBS + 20 mM Cysteine-HCL+ 10 mM EDTA, pH = 7.4) for 5 h at 37° C. We stopped the reaction by centrifuging the mix in a Spin-X centrifuge and removed residual IgG1 and Fc fragments by incubating the flow-through with protein A agarose for 2 h at room temperature. Flow-through was injected into a Superdex200 column and peak fractions corresponding to IGH505 Fab were collected and concentrated in Vivaspin6 10-kDa filter units. The polyclonal IgG serum pool used in Figure 1a consist of serum samples from four HCV^+^HIV^+^ individuals of the MOSAIC cohort. The two serum pools were diluted 1:4 in PBS before purification with protein G or protein A agarose resin (Pierce) according to manufacturer’s instruction. The eluted polyclonal IgG serum was then concentrated with a Vivaspin6 10-kDa cutoff filters, volume was adjusted to match the input serum volume using PBS and stored at 4° C.

### Neutralization assays

To generate pseudoparticles (HCVpp), 293T or 293T CD81KO cells were co-transfected with three plasmids: a MLV (phCMV-5349) packaging construct, a luciferase reporter plasmid and an expression vector encoding the full-length H77 or AMS0232 E1E2 glycoprotein complex^58^. Supernatants containing HCVpp were collected after 72 h and stored at −80° C until further use. For neutralization studies, pseudovirus-containing supernatant was incubated in duplicates or quadruplicates with a dilution series of purified polyclonal human IgG or mAbs prior to infection. Virus-antibody mixtures were incubated for 1 h at 37 ° C and were added to Huh-7 cells and incubated for 4h before addition of DMEM culture media. Infection was allowed to proceed for 72 h. For readout, the samples were lysed and assayed using Firefly luciferase and a GloMax luminometer (Promega, USA). Data was analysed in GraphPad Prism 8.0.

### ELISA

Purified E1E2 + AR4A-StrepII complexes (1.0 μg ml^™1^ in TBS with 0.1% DDM) were coated for 2 h at room temperature on 96-well Strep-Tactin XT coated microplates (IBA LifeSciences). Plates were washed twice with TBS/0.05% Tween20 before incubating with serially diluted mAbs and CD81-LEL-hFc (R&D systems) in casein blocking buffer (Thermo Fisher Scientific) diluted 1:1 v:v in TBS with 0.1% DDM (casein/0.1% DDM) for 90 min. After three washes with TBS/0.05% Tween20, HRP-labelled goat anti-human IgG (Jackson Immunoresearch) (1:3000) in casein/0.1% DDM was added for 45 min. After five washes with TBS/0.05% Tween20, plates were developed by adding develop solution [1% 3,3′,5,5′-tetraethylbenzidine (Sigma-Aldrich), 0.01 % H_2_O_2_, 100 mM sodium acetate, 100 mM citric acid] and the reaction was stopped after 5 min by adding 0.8 M H_2_SO_4_. Absorbance was measured at 450 nm and data was visualized in GraphPad Prism.

### Purification of E1E2 complexes

HEK293F cells were transfected at a density of ∼1.5 × 10^6^ cells/ml by mixing PEIMAX with plasmids containing full-length AMS0232 E1E2, StrepII-tagged AR4A HC Fab and AR4A LC in a 1:1.3:1.9 ratio. Cell pellets were harvested 72 h post transfection, washed with ice-cold PBS and solubilized in PBS containing 1% triton-X-100 and cOmplete protease inhibitor cocktail tablets (Sigma Aldrich) (1.5 tablets/L of transfected cells). The lysate was incubated at 4 ° C for 30 min, centrifuged at 4,000g for 30 min, filtered and flowed over a Strep-Tactin XT 4Flow affinity resin. Resin was washed with twelve resin volumes of wash buffer 1 (0.1 M Tris-Cl, 0.15 M NaCl, 1 mM EDTA, 0.025% triton-X-100, pH 8) and sixteen resin volumes of wash buffer 2 (0.1 M Tris-Cl, 0.15 M NaCl, 1 mM EDTA, 0.1% *n*-dodecyl β-D-maltoside (DDM), pH 8). The complexes were eluted with buffer BXT supplemented with DDM (0.1 M Tris-Cl, 0.15 M NaCl, 1 mM EDTA, 50 mM biotin, 0.1% DDM, pH 8). The eluted proteins were concentrated in 100-kDa cutoff filters and then injected into a Superose 6 increase column equilibrated in TBS/0.1% DDM.

### Sample preparation for cryo-EM

Five plasmids were co-transfected (E1E2, AR4A heavy chain, AR4A light chain, AT12009 heavy chain and AT12009 light chain at a ratio of 1:1.2:1.5:1.7:2, respectively). Cell pellets were harvested 72 h post transfection, washed with ice-cold PBS and solubilized in PBS containing 1% triton-X-100 and cOmplete protease inhibitor cocktail tablets (Sigma Aldrich) (1.5 tablets/L of transfected cells). The lysate was incubated at 4 ° C for 1 h, and clarified by centrifugation in a JLA 16.25 rotor at 39,500g for 1 h. The clarified lysate was flowed over Streptactin XT 4Flow, washed but before elution, the resin was first resuspended in 2ml of wash buffer two and 500x molar excess of peptidisc (peptidisc biotech) was added and incubated at 4 ° C for 1 h. 100 mg of biobeads SM-2 (BioRad) were added to the resin containing the protein of interest and the peptidiscs and incubated O/N at 4 ° C. Flowthrough containing excess of peptidisc excess was discarded and five resin volumes of elution buffer (0.1 M Tris-Cl, 0.15 M NaCl, 1 mM EDTA, 50 mM biotin, pH 8) were added to the resin. The protein was eluted slowly (6-24 h), then incubated with 5x molar excess of IGH505 Fab for 1 h at 4°C. The complex was concentrated in a 30-kDa cutoff centrifugal filter unit and then injected into a Superose 6 increase column in TBS. Peak fractions corresponding to E1E2 in complex with AR4A, AT12009 and IGH505 Fabs were concentrated to 1mg ml^-1^. Protein was measured by absorbance at 280 nm using 1 ABS = 1 mg ml^-1^.

Next, UltrAuFoil R 1.2/1.3 300 mesh gold grids were plasma cleaned for 7 s using a Solarus advanced plasma cleaning system (Gatan) before loading the sample (Ar/O_2_; 35 sccm; 50W). Subsequently, 3 μl of purified protein was loaded onto the grid and plunge-frozen into nitrogen-cooled liquid ethane using the Vitrobot mark IV (Thermo Fisher Scientific). The settings for the Vitrobot were as follows: Temperature: 4 ° C, humidity: 100%, blotting time: 5 s with standard Vitrobot filter paper, Ø55/20mm, Grade 595, blotting force: 0 and wait time: 7 s.

### Cryo-EM data collection

Grids were loaded into a Talos Arctica transmission electron microscope (Thermo Fischer Scientific) operating at 200kV and images acquired using a K2 Summit direct electron detector camera (Gatan). The data were collected at a total cumulative exposure of 50e^-^/Å^2^, 9 s exposure time, 36 frames and 250 ms/frame (See Table 1 for more details). Magnification was set at 36,000x with a resulting pixel size of 1.15 Å/pix at the specimen plane. Automated data collection was performed using Leginon software^59^. The data collection details are presented in Table S1.

### Cryo-EM image processing

Image preprocessing was performed using the Appion image processing environment^60^. Stacks of dose-fractionated image frames were aligned using the UCSF MotionCor2 software^61^. Particles were selected from the drift-corrected images (initial dataset, 2,650 images; second dataset, 2,471 images; for a total of 5,121 micrographs) using the cryoSPARC image processing suite^62^. We estimated the CTF using GCTF^63^ and extracted our particles at 1.15 Å/pixel with a box size of 300 × 300 pixels. A total of 2,017,845 particles were extracted from micrographs and 2D classified^62^. A total of 1,049,598 particles were selected based on signal to noise ratio for further processing in 3D. Two rounds of heterogeneous classification (n=2 and n=4, respectively, with default parameters) produced one 3D class containing 34% of the particles that represented E1E2 in complex with three Fabs (AT12009 Fab, AR4A Fab and IGH505 Fab). Additionally, two classes containing 42% of the particles resembled E1E2 in complex with two Fabs (IGH505 Fab and AR4A Fab) (Supplementary Fig. 2C). In total, 355,643 particles from the representative class that we hypothesized contained E1E2 in complex with three Fabs was selected for non-uniform refinement, which generated a model with a reported a resolution at 3.49 Å based on an Fourier Shell Correlation (FSC) cutoff at 0.143. The 3.49 Å E1E2 reconstruction was further analyzed using 3D variability analysis (3DVA) using 3 eigenvectors and 8 clusters. Two clusters were selected based on visible secondary structural elements in E1 as well as a more defined peptidisc region. To ensure the density in this region was due to signal from E1 and not noise, the 48,546 particles from these two clusters were used for 3D non-uniform refinement. The resulting model resolved to 3.83Å and provided additional insight into the structure of E1, particularly its membrane associated helices. We masked the flexible regions of E1 (highlighted in Supplemental Fig. 2D) for local refinement using all 1,049,598 particles. Local refinement generated a 3.37 Å map that enabled us to resolve additional, highly flexible regions of E1.

### Model building and refinement

Two postprocessed maps were used to build the final atomic models. The first map contains E1, E2 and the 3 Fabs. Phenix map-to-model option was used to build E1 and E2 into the model^64^. Two fragments of 15 amino acids were chosen to start de novo building in both models. Precise sequence registry assignment was determined by locating bulky Phe, Arg, Tyr, and Trp side chains. Iterative manual model building and automatic refinement were carried out in Coot. ABodyBuilder was used to produce the models of both antibody Fv regions^65^. Multiple rounds of Rosetta relaxed refinement^41^ and manual Coot refinement were performed to build the final models. Model fit to map was validated using MolProbity^66^ and EMRinger^67^ analyses. The final refined models were submitted to the Protein Data Bank (PDB). All structural figures were generated using UCSF ChimeraX^68^.

### AlphaFold Prediction Software

AlphaFold was used as a prediction software to determine the atomic model for both E1 and E2 individually. Specifically, the jupyter notebook module of ColabFold (AlphaFold2_advanced.ipynb) was utilized and accessed from the following link: https://colab.research.google.com/github/sokrypton/ColabFold/blob/main/beta/AlphaFold2_advanced.ipynb#scrollTo=pc5-mbsX9PZC

The run parameters were as followed: (1) sequence (for E1 and E2, respectively): “YQVRNSTGLYHVTNDCPNSSIVYETADAILHTPGCVPCVREGNASRCWVPMTPTVATRDGKLPATQLRRHIDLLVGSATLCSALYVGDLCGSVFLVGQLFTFSPRRHWTTQDCNCSIYPGHVTGHRMAWDMMMNWSPTTALVVAQLLRIPQAILDMIAGAHWGVLAGLAYFSMVGNWAKVLAVLLLFAGVDA”, “QTYVTGGTAARATSGLANFFSPGAKQDVQLINTNGSWHINRTALNCNTSLETGWIAGLIYLNKFNSSGCPERMASCRPLADFAQGWGPISYANGSGPDHRPYCWHYPPKPCGIVSAKSVCGPVYCFTPSPVVVGTTNKLGAPTYSWGENETDVFVLNNTRPPLGNWFGCTWMNSTGFTKVCGAPPCAIGGVGNNTLHCPTDCFRKHPEATYSRCGSGPWITPRCLVDYPYRLWHYPCTINYTRFKVRMYIGGVEHRLDAACNWTRGERCDLEDRDRSELSPLLLSTTQWQVLPCSFTTLPALSTGLIHLHQNIVDVQYLYGVGSSIVSWAIKWEYVVLLFLLLADARVCSCLWMMLLISQAEA” (2) jobname: “E1” or “E2” and (3) homooligomer: 1.

### Site-specific glycan analysis

Two aliquots of the purified E1E2+AR4A+AT12009 Fab sample were denatured for 1h in 50 mM Tris/HCl, pH 8.0 containing 6 M of urea and 5 mM dithiothreitol (DTT). Next, the sample was reduced and alkylated by adding 20 mM iodoacetamide (IAA) and incubated for 1h in the dark, followed by a 1h incubation with 20 mM DTT to eliminate residual IAA. The alkylated E1E2 proteins were buffer-exchanged into 50 mM Tris/HCl, pH 8.0 using Vivaspin columns (3 kDa) and two of the aliquots were digested separately overnight using chymotrypsin (Mass Spectrometry Grade, Promega) or alpha lytic protease (Sigma Aldrich) at a ratio of 1:30 (w/w). The next day, the peptides were dried and extracted using C18 Zip-tip (MerckMilipore). The peptides were dried again, re-suspended in 0.1% formic acid and analyzed by nanoLC-ESI MS with an Ultimate 3000 HPLC (Thermo Fisher Scientific) system coupled to an Orbitrap Eclipse mass spectrometer (Thermo Fisher Scientific) using stepped higher energy collision-induced dissociation (HCD) fragmentation. Peptides were separated using an EasySpray PepMap RSLC C18 column (75 µm × 75 cm). A trapping column (PepMap 100 C18 3μM, 75μM × 2cm) was used in line with the LC prior to separation with the analytical column. The LC conditions were as follows: 280 min linear gradient consisting of 4-32% acetonitrile in 0.1% formic acid over 260 minutes followed by 20 min of alternating 76% acetonitrile in 0.1% formic acid and 4% ACN in 0.1% formic acid, used to ensure all the sample had eluted from the column. The flow rate was set to 300 nL/min. The spray voltage was set to 2.5 kV and the temperature of the heated capillary was set to 40 °C. The ion transfer tube temperature was set to 275 °C. The scan range was 375−1500 m/z. Stepped HCD collision energy was set to 15, 25 and 45% and the MS2 for each energy was combined. Precursor and fragment detection were performed using an Orbitrap at a resolution MS1= 120,000. MS2= 30,000. The AGC target for MS1 was set to standard and injection time set to auto which involves the system setting the two parameters to maximize sensitivity while maintaining cycle time. Full LC and MS methodology can be extracted from the appropriate Raw file using XCalibur FreeStyle software or upon request. Glycopeptide fragmentation data were extracted from the raw file using Byos (Version 4.0; Protein Metrics Inc.). The glycopeptide fragmentation data were evaluated manually for each glycopeptide; the peptide was scored as true-positive when the correct b and y fragment ions were observed along with oxonium ions corresponding to the glycan identified. The MS data was searched using the Protein Metrics 305 N-glycan library with sulfated glycans added manually. To search for N-glycans at non-canonical motifs, the above library was instead added as a custom modification to search for glycans (common1) at any N. The relative amounts of each glycan at each site as well as the unoccupied proportion were determined by comparing the extracted chromatographic areas for different glycotypes with an identical peptide sequence. All charge states for a single glycopeptide were summed. The precursor mass tolerance was set at 4 ppm and 10 ppm for fragments. A 1% false discovery rate (FDR) was applied. The relative amounts of each glycan at each site as well as the unoccupied proportion were determined by comparing the extracted ion chromatographic areas for different glycopeptides with an identical peptide sequence.

Glycan compositions were grouped according to their level of processing. HexNAc(2)Hex(10–5) compositions were classified as oligomannose-type, HexNAc(3)Hex(5-6)/HexNac(3)Hex(5-6)Fuc were classified as hybrid, and remaining glycan compositions were classified as complex-type.

## Data availability

The PDBs and EMDBs for E1E2 maps have been deposited into the RCSB PDB (https://www.rcsb.org) under accession number 7T6X and to the EMDB database (https://www.ebi.ac.uk/emdb/) under the accession number EMD-25730. The mass spectrometry RAW files have been deposited in the MASSive server (https://massive.ucsd.edu) with accession number (MSV000088553).

## References

1 Fuerst, T. R., Pierce, B. G., Keck, Z. Y. & Foung, S. K. H. Designing a B Cell-Based Vaccine against a Highly Variable Hepatitis C Virus. Front Microbiol 8, 2692, doi:10.3389/fmicb.2017.02692 (2017).

2 Yost, S. A., Wang, Y. & Marcotrigiano, J. Hepatitis C Virus Envelope Glycoproteins: A Balancing Act of Order and Disorder. Front Immunol 9, 1917, doi:10.3389/fimmu.2018.01917 (2018).

3 Kinchen, V. J. et al. Broadly Neutralizing Antibody Mediated Clearance of Human Hepatitis C Virus Infection. Cell Host Microbe 24, 717–730 e715, doi:10.1016/j.chom.2018.10.012 (2018).

4 Pestka, J. M. et al. Rapid induction of virus-neutralizing antibodies and viral clearance in a single-source outbreak of hepatitis C. Proc Natl Acad Sci U S A 104, 6025–6030, doi:10.1073/pnas.0607026104 (2007).

5 Merat, S. J. et al. Cross-genotype AR3-specific neutralizing antibodies confer long-term protection in injecting drug users after HCV clearance. J Hepatol 71, 14–24, doi:10.1016/j.jhep.2019.02.013 (2019).

6 Law, M. et al. Broadly neutralizing antibodies protect against hepatitis C virus quasispecies challenge. Nat Med 14, 25–27, doi:10.1038/nm1698 (2008).

7 O’Shea, D. et al. Prevention of hepatitis C virus infection using a broad cross-neutralizing monoclonal antibody (AR4A) and epigallocatechin gallate. Liver Transpl 22, 324–332, doi:10.1002/lt.24344 (2016).

8 de Jong, Y. P. et al. Broadly neutralizing antibodies abrogate established hepatitis C virus infection. Sci Transl Med 6, 254ra129, doi:10.1126/scitranslmed.3009512 (2014).

9 Lindenbach, B. D. & Rice, C. M. The ins and outs of hepatitis C virus entry and assembly. Nat Rev Microbiol 11, 688–700, doi:10.1038/nrmicro3098 (2013).

10 Pileri, P. et al. Binding of hepatitis C virus to CD81. Science 282, 938–941, doi:10.1126/science.282.5390.938 (1998).

11 Scarselli, E. et al. The human scavenger receptor class B type I is a novel candidate receptor for the hepatitis C virus. Embo j 21, 5017–5025, doi:10.1093/emboj/cdf529 (2002).

12 Tong, M. J., Theodoro, C. F. & Salvo, R. T. Late development of hepatocellular carcinoma after viral clearance in patients with chronic hepatitis C: A need for continual surveillance. J Dig Dis 19, 411–420, doi:10.1111/1751-2980.12615 (2018).

13 Drummer, H. E., Boo, I. & Poumbourios, P. Mutagenesis of a conserved fusion peptide-like motif and membrane-proximal heptad-repeat region of hepatitis C virus glycoprotein E1. J Gen Virol 88, 1144–1148, doi:10.1099/vir.0.82567-0 (2007).

14 Catanese, M. T. et al. Ultrastructural analysis of hepatitis C virus particles. Proc Natl Acad Sci U S A 110, 9505–9510, doi:10.1073/pnas.1307527110 (2013).

15 Vieyres, G., Dubuisson, J. & Pietschmann, T. Incorporation of hepatitis C virus E1 and E2 glycoproteins: the keystones on a peculiar virion. Viruses 6, 1149–1187, doi:10.3390/v6031149 (2014).

16 Stejskal, L. et al. Flexibility and intrinsic disorder are conserved features of hepatitis C virus E2 glycoprotein. PLoS Comput Biol 16, e1007710, doi:10.1371/journal.pcbi.1007710 (2020).

17 Lavie, M., Hanoulle, X. & Dubuisson, J. Glycan Shielding and Modulation of Hepatitis C Virus Neutralizing Antibodies. Front Immunol 9, 910, doi:10.3389/fimmu.2018.00910 (2018).

18 Goffard, A. et al. Role of N-linked glycans in the functions of hepatitis C virus envelope glycoproteins. J Virol 79, 8400–8409, doi:10.1128/JVI.79.13.8400-8409.2005 (2005).

19 Prentoe, J. et al. Hypervariable region 1 and N-linked glycans of hepatitis C regulate virion neutralization by modulating envelope conformations. Proc Natl Acad Sci U S A 116, 10039–10047, doi:10.1073/pnas.1822002116 (2019).

20 Kong, L. et al. Hepatitis C virus E2 envelope glycoprotein core structure. Science 342, 1090–1094, doi:10.1126/science.1243876 (2013).

21 Khan, I. et al. Modulation of hepatitis C virus genome replication by glycosphingolipids and four-phosphate adaptor protein 2. J Virol 88, 12276–12295, doi:10.1128/JVI.00970-14 (2014).

22 Flyak, A. I. et al. HCV Broadly Neutralizing Antibodies Use a CDRH3 Disulfide Motif to Recognize an E2 Glycoprotein Site that Can Be Targeted for Vaccine Design. Cell Host Microbe 24, 703–716.e703, doi:10.1016/j.chom.2018.10.009 (2018).

23 Tzarum, N. et al. Genetic and structural insights into broad neutralization of hepatitis C virus by human VH1-69 antibodies. Sci Adv 5, eaav1882, doi:10.1126/sciadv.aav1882 (2019).

24 Tzarum, N., Wilson, I. A. & Law, M. The Neutralizing Face of Hepatitis C Virus E2 Envelope Glycoprotein. Front Immunol 9, 1315, doi:10.3389/fimmu.2018.01315 (2018).

25 Kumar, A. et al. Structural insights into hepatitis C virus receptor binding and entry. Nature 598, 521–525, doi:10.1038/s41586-021-03913-5 (2021).

26 El Omari, K. et al. Unexpected structure for the N-terminal domain of hepatitis C virus envelope glycoprotein E1. Nat Commun 5, 4874, doi:10.1038/ncomms5874 (2014).

27 Potter, J. A. et al. Toward a hepatitis C virus vaccine: the structural basis of hepatitis C virus neutralization by AP33, a broadly neutralizing antibody. J Virol 86, 12923–12932, doi:10.1128/JVI.02052-12 (2012).

28 Kong, L. et al. Structure of Hepatitis C Virus Envelope Glycoprotein E1 Antigenic Site 314-324 in Complex with Antibody IGH526. J Mol Biol 427, 2617–2628, doi:10.1016/j.jmb.2015.06.012 (2015).

29 Giang, E. et al. Human broadly neutralizing antibodies to the envelope glycoprotein complex of hepatitis C virus. Proc Natl Acad Sci U S A 109, 6205–6210, doi:10.1073/pnas.1114927109 (2012).

30 Ruwona, T. B., Giang, E., Nieusma, T. & Law, M. Fine mapping of murine antibody responses to immunization with a novel soluble form of hepatitis C virus envelope glycoprotein complex. J Virol 88, 10459–10471, doi:10.1128/JVI.01584-14 (2014).

31 Cao, L. et al. Functional expression and characterization of the envelope glycoprotein E1E2 heterodimer of hepatitis C virus. PLoS Pathog 15, e1007759, doi:10.1371/journal.ppat.1007759 (2019).

32 Guest, J. D. et al. Design of a native-like secreted form of the hepatitis C virus E1E2 heterodimer. Proc Natl Acad Sci U S A 118, doi:10.1073/pnas.2015149118 (2021).

33 Thomas, X. V. et al. Genetic characterization of multiple hepatitis C virus infections following acute infection in HIV-infected men who have sex with men. AIDS 29, 2287–2295, doi:10.1097/QAD.0000000000000838 (2015).

34 Carlson, M. L. et al. The Peptidisc, a simple method for stabilizing membrane proteins in detergent-free solution. Elife 7, doi:10.7554/eLife.34085 (2018).

35 Schaefer, W. et al. Immunoglobulin domain crossover as a generic approach for the production of bispecific IgG antibodies. Proc Natl Acad Sci U S A 108, 11187–11192, doi:10.1073/pnas.1019002108 (2011).

36 Punjani, A. & Fleet, D. J. 3D variability analysis: Resolving continuous flexibility and discrete heterogeneity from single particle cryo-EM. J Struct Biol 213, 107702, doi:10.1016/j.jsb.2021.107702 (2021).

37 Castelli, M. et al. HCV E2 core structures and mAbs: something is still missing. Drug Discov Today 19, 1964–1970, doi:10.1016/j.drudis.2014.08.011 (2014).

38 Wahid, A. et al. Disulfide bonds in hepatitis C virus glycoprotein E1 control the assembly and entry functions of E2 glycoprotein. J Virol 87, 1605–1617, doi:10.1128/JVI.02659-12 (2013).

39 Krey, T. et al. The disulfide bonds in glycoprotein E2 of hepatitis C virus reveal the tertiary organization of the molecule. PLoS Pathog 6, e1000762, doi:10.1371/journal.ppat.1000762 (2010).

40 Marin, M. Q. et al. Optimized Hepatitis C Virus (HCV) E2 Glycoproteins and their Immunogenicity in Combination with MVA-HCV. Vaccines (Basel) 8, doi:10.3390/vaccines8030440 (2020).

41 Wang, W. et al. Alanine scanning mutagenesis of hepatitis C virus E2 cysteine residues: Insights into E2 biogenesis and antigenicity. Virology 448, 229–237, doi:10.1016/j.virol.2013.10.020 (2014).

42 Deleersnyder, V. et al. Formation of native hepatitis C virus glycoprotein complexes. J Virol 71, 697–704, doi:10.1128/JVI.71.1.697-704.1997 (1997).

43 Di Lorenzo, C., Angus, A. G. & Patel, A. H. Hepatitis C virus evasion mechanisms from neutralizing antibodies. Viruses 3, 2280–2300, doi:10.3390/v3112280 (2011).

44 Meunier, J. C. et al. Analysis of the glycosylation sites of hepatitis C virus (HCV) glycoprotein E1 and the influence of E1 glycans on the formation of the HCV glycoprotein complex. J Gen Virol 80 (Pt 4), 887–896, doi:10.1099/0022-1317-80-4-887 (1999).

45 Vieyres, G. et al. Characterization of the envelope glycoproteins associated with infectious hepatitis C virus. J Virol 84, 10159–10168, doi:10.1128/JVI.01180-10 (2010).

46 Cocquerel, L. et al. The transmembrane domain of hepatitis C virus glycoprotein E1 is a signal for static retention in the endoplasmic reticulum. J Virol 73, 2641–2649, doi:10.1128/JVI.73.4.2641-2649.1999 (1999).

47 Gopal, R. et al. Probing the antigenicity of hepatitis C virus envelope glycoprotein complex by high-throughput mutagenesis. PLoS Pathog 13, e1006735, doi:10.1371/journal.ppat.1006735 (2017).

48 Meunier, J. C. et al. Isolation and characterization of broadly neutralizing human monoclonal antibodies to the e1 glycoprotein of hepatitis C virus. J Virol 82, 966–973, doi:10.1128/JVI.01872-07 (2008).

49 Michalak, J. P. et al. Characterization of truncated forms of hepatitis C virus glycoproteins. J Gen Virol 78 (Pt 9), 2299–2306, doi:10.1099/0022-1317-78-9-2299 (1997).

50 Patel, J., Patel, A. H. & McLauchlan, J. The transmembrane domain of the hepatitis C virus E2 glycoprotein is required for correct folding of the E1 glycoprotein and native complex formation. Virology 279, 58–68, doi:10.1006/viro.2000.0693 (2001).

51 Brazzoli, M. et al. Folding and dimerization of hepatitis C virus E1 and E2 glycoproteins in stably transfected CHO cells. Virology 332, 438–453, doi:10.1016/j.virol.2004.11.034 (2005).

52 McLellan, J. S. et al. Structure of RSV fusion glycoprotein trimer bound to a prefusion-specific neutralizing antibody. Science 340, 1113–1117, doi:10.1126/science.1234914 (2013).

53 de Taeye, S. W. et al. Immunogenicity of Stabilized HIV-1 Envelope Trimers with Reduced Exposure of Non-neutralizing Epitopes. Cell 163, 1702–1715, doi:10.1016/j.cell.2015.11.056 (2015).

54 Juraszek, J. et al. Stabilizing the closed SARS-CoV-2 spike trimer. Nat Commun 12, 244, doi:10.1038/s41467-020-20321-x (2021).

55 Falson, P. et al. Hepatitis C Virus Envelope Glycoprotein E1 Forms Trimers at the Surface of the Virion. J Virol 89, 10333–10346, doi:10.1128/JVI.00991-15 (2015).

56 Emsley, P. & Crispin, M. Structural analysis of glycoproteins: building N-linked glycans with Coot. Acta Crystallogr D Struct Biol 74, 256–263, doi:10.1107/S2059798318005119 (2018).

57 Kuiken, C. et al. Hepatitis C databases, principles and utility to researchers. Hepatology 43, 1157–1165, doi:10.1002/hep.21162 (2006).

58 Kalemera, M. D. et al. Optimized cell systems for the investigation of hepatitis C virus E1E2 glycoproteins. J Gen Virol 102, doi:10.1099/jgv.0.001512 (2021).

59 Suloway, C. et al. Automated molecular microscopy: the new Leginon system. J Struct Biol 151, 41–60, doi:10.1016/j.jsb.2005.03.010 (2005).

60 Lander, G. C. et al. Appion: an integrated, database-driven pipeline to facilitate EM image processing. J Struct Biol 166, 95–102, doi:10.1016/j.jsb.2009.01.002 (2009).

61 Zheng, S. Q. et al. MotionCor2: anisotropic correction of beam-induced motion for improved cryo-electron microscopy. Nat Methods 14, 331–332, doi:10.1038/nmeth.4193 (2017).

62 Punjani, A., Rubinstein, J. L., Fleet, D. J. & Brubaker, M. A. cryoSPARC: algorithms for rapid unsupervised cryo-EM structure determination. Nat Methods 14, 290–296, doi:10.1038/nmeth.4169 (2017).

63 Zhang, K. Gctf: Real-time CTF determination and correction. J Struct Biol 193, 1–12, doi:10.1016/j.jsb.2015.11.003 (2016).

64 Liebschner, D. et al. Macromolecular structure determination using X-rays, neutrons and electrons: recent developments in Phenix. Acta Crystallogr D Struct Biol 75, 861–877, doi:10.1107/S2059798319011471 (2019).

65 Leem, J., Dunbar, J., Georges, G., Shi, J. & Deane, C. M. ABodyBuilder: Automated antibody structure prediction with data-driven accuracy estimation. MAbs 8, 1259–1268, doi:10.1080/19420862.2016.1205773 (2016).

66 Williams, C. J. et al. MolProbity: More and better reference data for improved all-atom structure validation. Protein Sci 27, 293–315, doi:10.1002/pro.3330 (2018).

67 Barad, B. A. et al. EMRinger: side chain-directed model and map validation for 3D cryo-electron microscopy. Nat Methods 12, 943–946, doi:10.1038/nmeth.3541 (2015).

68 Pettersen, E. F. et al. UCSF ChimeraX: Structure visualization for researchers, educators, and developers. Protein Sci 30, 70–82, doi:10.1002/pro.3943 (2021).

