## Supplemental Materials for "Structure of the hepatitis C virus E1E2 glycoprotein complex"

**Supplementary Table 1. Data collection, image processing and map and model refinement parameters.**

| | AMS0232-IGH505-AR4-AT12009 (EMDB 25730, PDB 7T6X) | C-terminal domain of E1-AMS0232 |
|---|---|---|
| **Data collection** | | |
| Microscope | Talos Arctica | |
| Voltage (keV) | 200 | |
| Detector | K2 Summit | |
| Magnification (nominal/calibrated) | 36,000X/43,478X | |
| Exposure navigation * | Image shift 4/16 holes | |
| Data acquisition software | Leginon | |
| Total electron exposure (e-/Å$^2$) | 50 | |
| Exposure rate (e-/pixel/sec) * | 6.074/6.754 | |
| Frame length (ms) | 250 | |
| No. of frames per micrograph | 49 | |
| Pixel size (Å) | 1.15 | |
| Defocus range (μm) | -0.8 to -1.12 | |
| Micrographs collected (no.) | 5,121 | |
| **Reconstruction** | | |
| Image processing package | Cryosparc | |
| Total extracted particles (no.) | 1049598 | |
| Refined particles (no.) | 355,643 | 993,212 |
| Final particles (no.) | 48,546 | 167,662 |
| Symmetry imposed | C1 | C1 |
| **Resolution (Å)** | | |
| <u>Map to Map</u> | | |
| FSC 0.5 (unmasked/loose masked/ tight masked) | 8.1/4.9/4.2 | 6.8/5.2/4.3 |
| FSC 0.143 (unmasked/loose masked/ tight masked) | 4.6/4.1/3.8 | 4.8/3.8/3.7 |
| Resolution range (local) | 3.0 - 4.5 | 3.0 - 10.0 |
| 3DFSC Sphericity | 0.949 out of 1 | 0.915 out of 1 |
| Sharpening B-factor (Å$^2$) | -109 | -117 |
| **Model Composition** | | |
| Protein residues | 1,140 | |
| Ligands | 16 | |
| N-acetyl-β-D-glucoseamine (NAG) | 24 | |
| α-D-mannose (MAN) | 6 | |
| β-D-mannose (BMA) | 7 | |
| **Model Refinement** | | |
| Refinement package | Phenix | |
| CC (volume / mask) | 0.82 / 0.83 | |

| | |
|---|---|
| R.m.s. deviations | |
| Bond lengths | 0.021 |
| Bond angles (°) | 1.942 |
| **Validation** | |
| Map-to-model FSC (0.5) | 3.9 |
| Ramachandran (%) | |
| Outliers | 0.09 |
| Allowed | 3.83 |
| Favored | 96.08 |
| MolProbity score | 1.33 |
| Poor rotamers (%) | 0 |
| Clashscore (all atoms) | 2.73 |
| C-beta deviations | 0 |
| CaBLAM Outliers (%) | 4.62 |
| EMRinger Score | 2.79 |

*These parameters include statistics that represent the following two datasets: 20nov20 and 21jul28.

**Supplementary Table 2. Buried surface area between E1E2 and the CDR loops of the AR4A, IGH505 and AT12009 bNAbs.**

| Antibody | Region | Interaction (subunit) | BSA (Å$^2$)* |
|---|---|---|---|
| **AR4A** | CDRH2 | E2 | 72 |
| | CDRH3 | E2 | 622 |
| **IGH505** | CDRH1 | E1 | 76 |
| | CDRH2 | E1 | 295 |
| | CDRH3 | E1 | 219 |
| | CDRL1 | E1 | 93 |
| | CDRL3 | E1 | 122 |
| | CDRL3 | E2 | 35 |
| **AT12009** | CDRH1 | E2 | 45 |
| | CDRH2 | E2 | 493 |
| | CDRH3 | E2 | 526 |

* Buried Surface Area (BSA) was calculated using PDBePISA

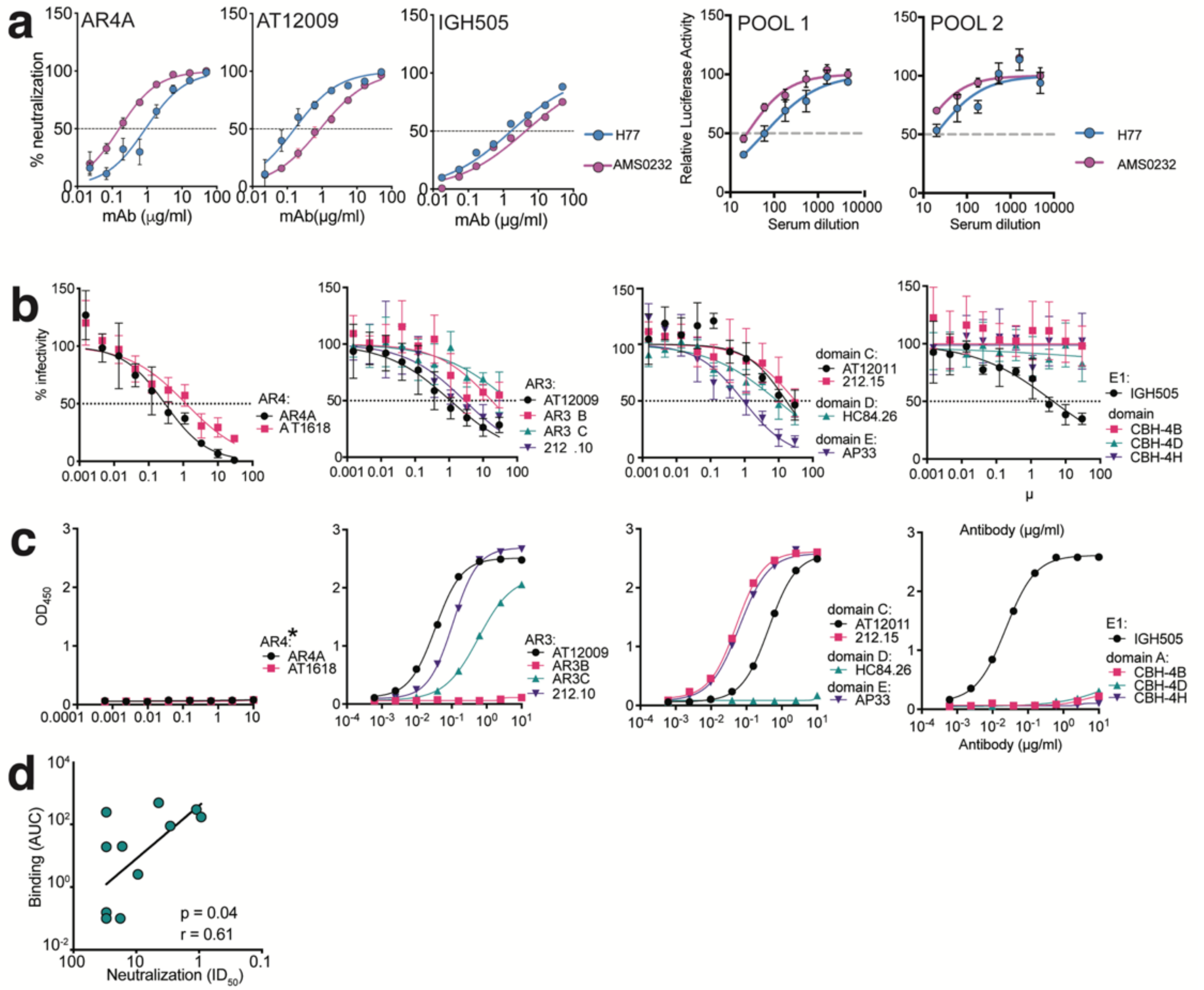

**Extended Data Fig. 1. Biochemical characterization of AMS0232 pseudovirus and full-length recombinant AMS0232 E1E2 glycoprotein complex**. (**a**) Pseudovirus neutralization experiments were performed in quadruplicates (n=4) for two pools of human sera and three monoclonal antibodies. Neutralization curves for bNAbs and serum pools to AMS0232 and H77 HCV pseudoviruses are plotted. (**b**) Neutralizing activity of monoclonal antibodies (tested in quadruplicate) against AMS0232 HCV pseudovirus. Error bars denote the S.D. Representative results from two independent experiments. (**c**) Binding of monoclonal antibodies to the purified E1E2 + AR4A Fab complex. Representative results from two independent experiments. (**d**) Correlation between neutralization ($ID_{50}$) from *b* and antibody binding (area under the curve (AUC)) from *c.* Values for AR4A and AT1618 were left out for this analysis. Spearman r and p-values are indicated

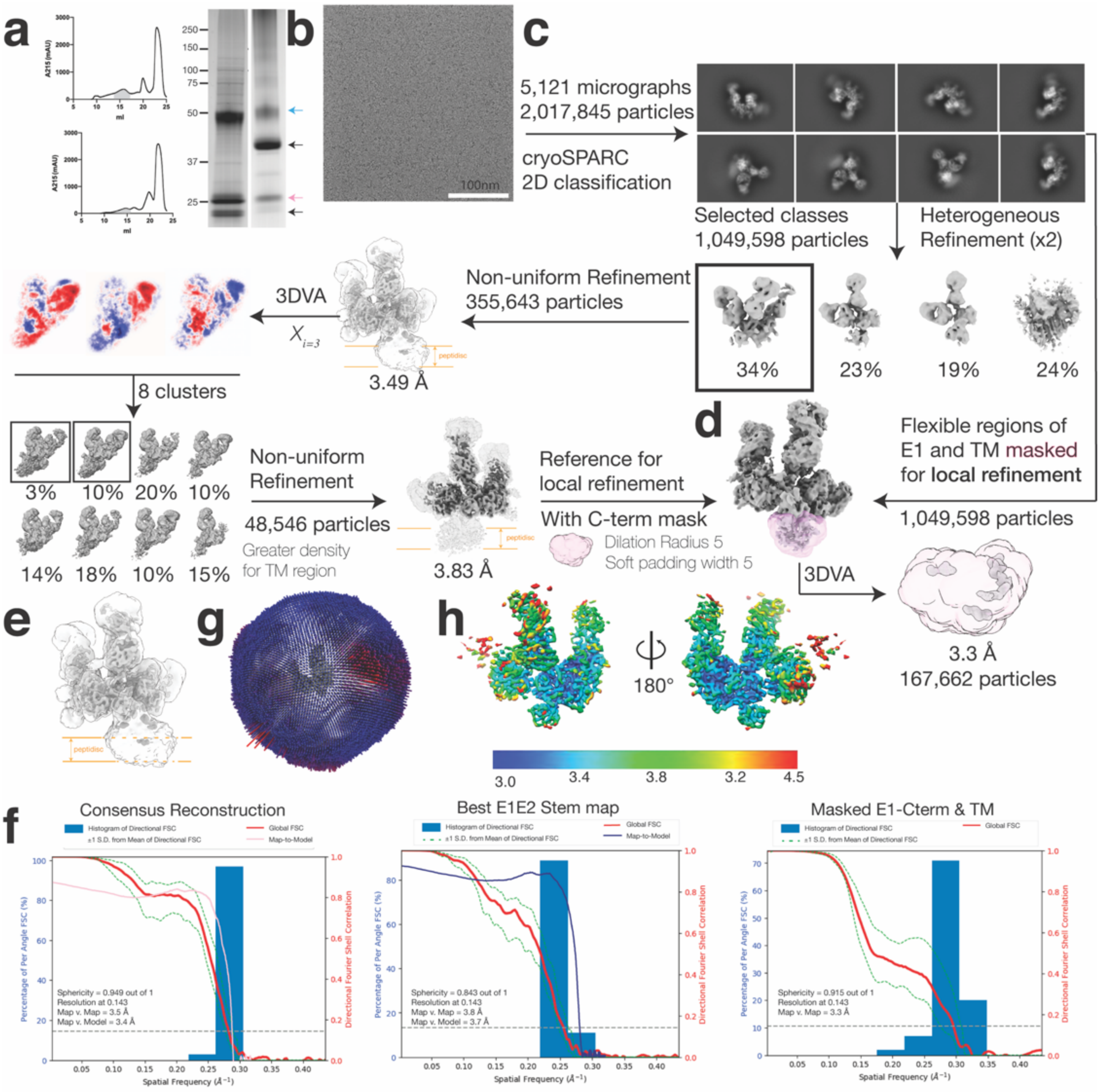

**Extended Data Fig. 2. CryoEM processing scheme of full-length HCV envelope glycoprotein E1E2 heterodimer in complex with bNAbs AT12009, IGH505 and AR4A.** (**a**) Size-exclusion chromatography curves for full-length envelope glycoprotein E1E2 heterodimer in complex with bNAbs AR4A and AT12009 embedded in a DDM micelle (top panel) and full-length envelope glycoprotein E1E2 heterodimer in complex with bNAbs AT12009, IGH505 and AR4A embedded in a peptidisc (bottom panel) are shown. The area corresponding to E1E2-bNAb complexes are highlighted in light gray. SDS-PAGE gels of E1E2 in complex with Fabs under reducing (left lane) and non-reducing (right lane) conditions, respectively. The blue arrow at 50 kDa represents E2, the pink arrow at 25 kDa represents E1, and the black arrows represent the Fabs. (**b**) Raw micrograph of full full-length HCV envelope glycoprotein E1E2 heterodimer in complex with bNAbs AT12009, IGH505, and AR4A collected on the Talos Arctica with a pixel size of 1.15 Å/pixel. (**c**) A total of 2,017,845 particles were extracted from micrographs and 2D/3Dclassified using the cryoSPARC image processing suite[57]. Next, 355,643 particles from the representative class that we hypothesized contained E1E2 in complex with three Fabs was selected for non-uniform refinement, which generated a model with a reported a resolution at 3.49Å. The final model resolved to 3.83Å and provided additional insight into the structure of E1, particularly its stem region. (**d**) Local refinement was performed to extract valuable signal from the stem and TMDs of E1E2. Using 1,049,598 particles from panel C, we ran multiple rounds of 3DVA and local refinement with the mask highlighted in pink to generate a 3.3 Å map of the E1 stem. The respective FSC curve of the local resolution map is to the right. (**e**) Overlay of the unsharpened 3.83 Å E1E2 map with threshold of 0.3 in ChimeraX with the sharpened 3.83 Å E1E2 map at a higher threshold of 0.1 in ChimeraX and the local resolution map to showcase the relative position of E1E2 to the membrane. (**f**) Three-Dimensional Fourier Shell Correlation (3DFSC) plot of the three maps. The $FSC_{map-map}$ resolution is calculated using 0.143 as the gold standard criterion, which represents how well the two half-maps from each dataset correlate as a function of spatial frequency. The $FSC_{map-model}$ curve, calculated in Phenix at a threshold of 0.143, was scaled proportionately and overlayed onto the following maps: (i) the consensus reconstruction and (ii) the Best E1E2 Stem map. (**g**) Angular distribution of the E1E2 maps. (**h**) Local resolution estimation for the consensus refinement.

### AMS0232 cryoEM structure

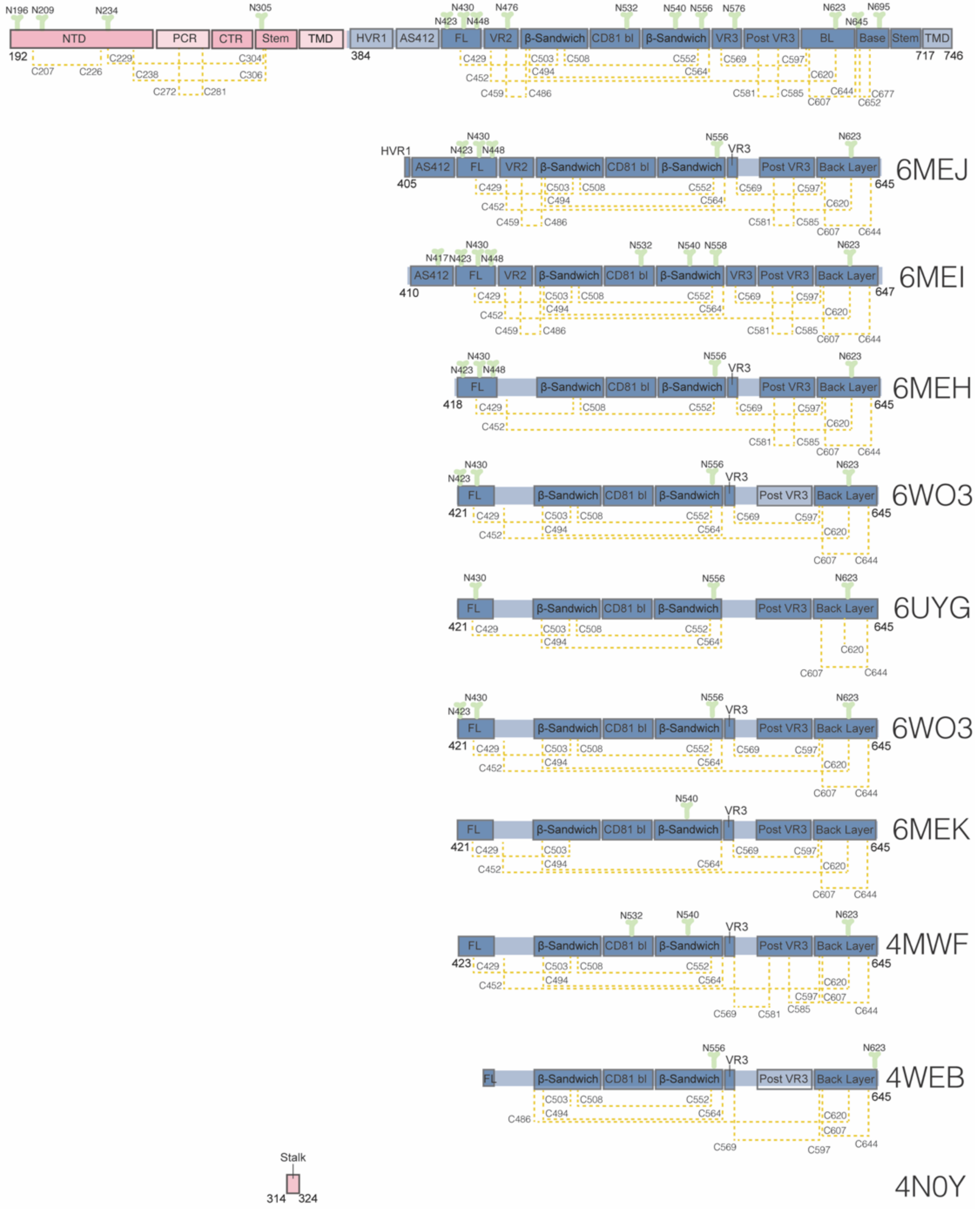

**Extended Data Fig. 3. Schematic representation of the regions resolved in our AMS0232 CryoEM structure compared to previous crystal structures of E1 and E2.** The solved regions are highlighted in dark pink and dark blue for E1 and E2, respectively. The unresolved regions are depicted in light pink and light blue. N-linked glycans are shown in green and numbered with their respective Asn residues. Disulfide bonds are shown in yellow dashed lines and numbered accordingly. PDB IDs from previous crystal structures are indicated.

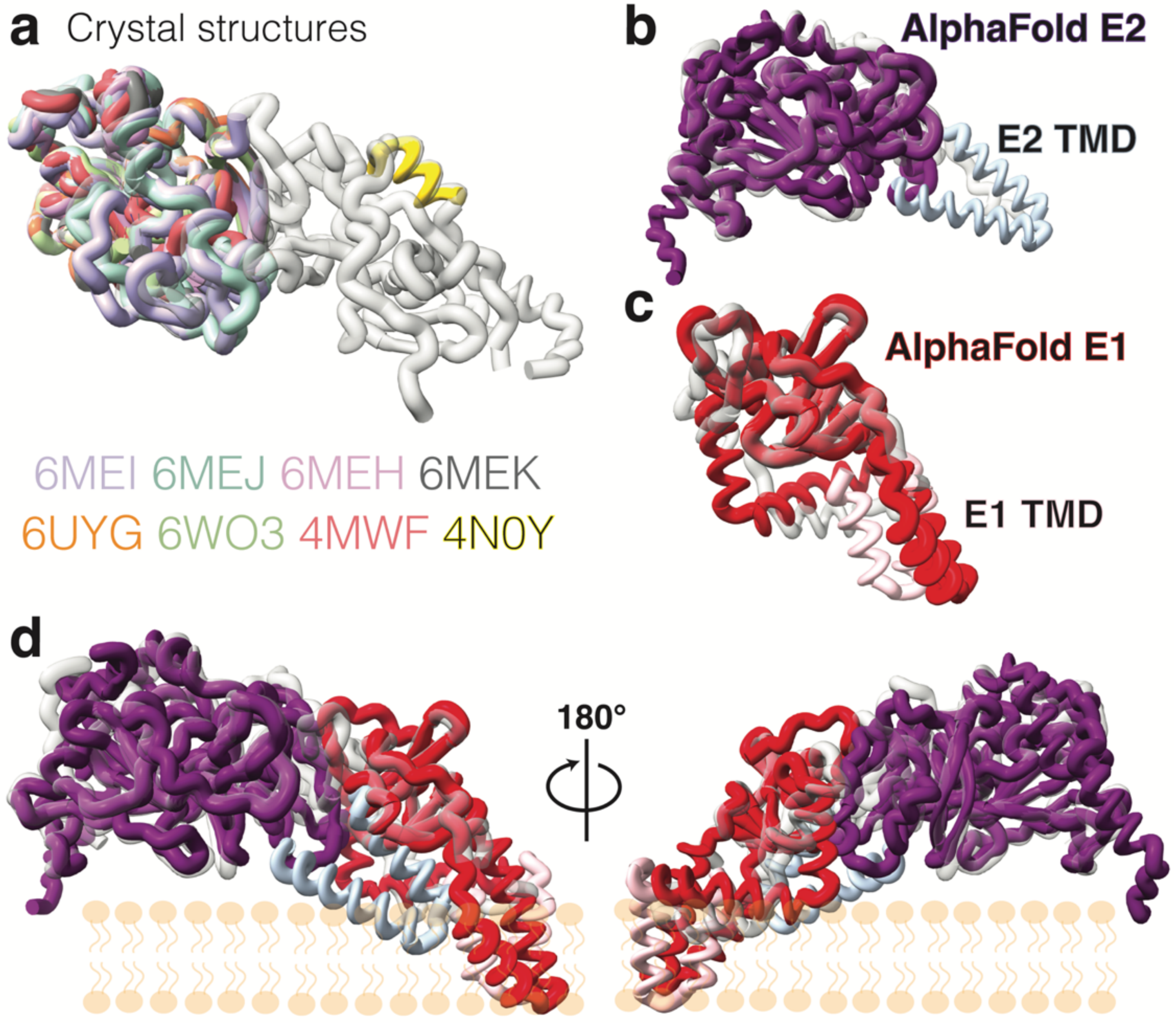

**Extended Data Fig. 4. Overlay of cryoEM full-length HCV E1E2 with published complementary crystal structures and the corresponding AlphaFold predictions.** (**a**) The following crystal structures of individual segments of E1 or E2 are superimposed to highlight the novel regions of E1E2 discovered by our cryoEM structure: E2, 6MEI, 6MEJ, 6MEH, 6MEK, 6UYG, 6WO3, 4MWF; E1, 4N0Y. Our cryoEM structure is transparent here for adequate comparison and further compared with the corresponding AlphaFold structures for (**b**) E2; in purple, with its TMD highlighted in light steel blue; and (**c**) E1; in red, with its TMD highlighted in pink. (**d**) Full-length E1E2 cryoEM structure (atomic model; spaghetti cartoon) superimposed with the AlphaFold E1 and E2 models. A lipid bilayer schematic represented in orange is shown to orient E1E2 heterodimer atomic model and the AlphaFold predicted model relative to the viral membrane.

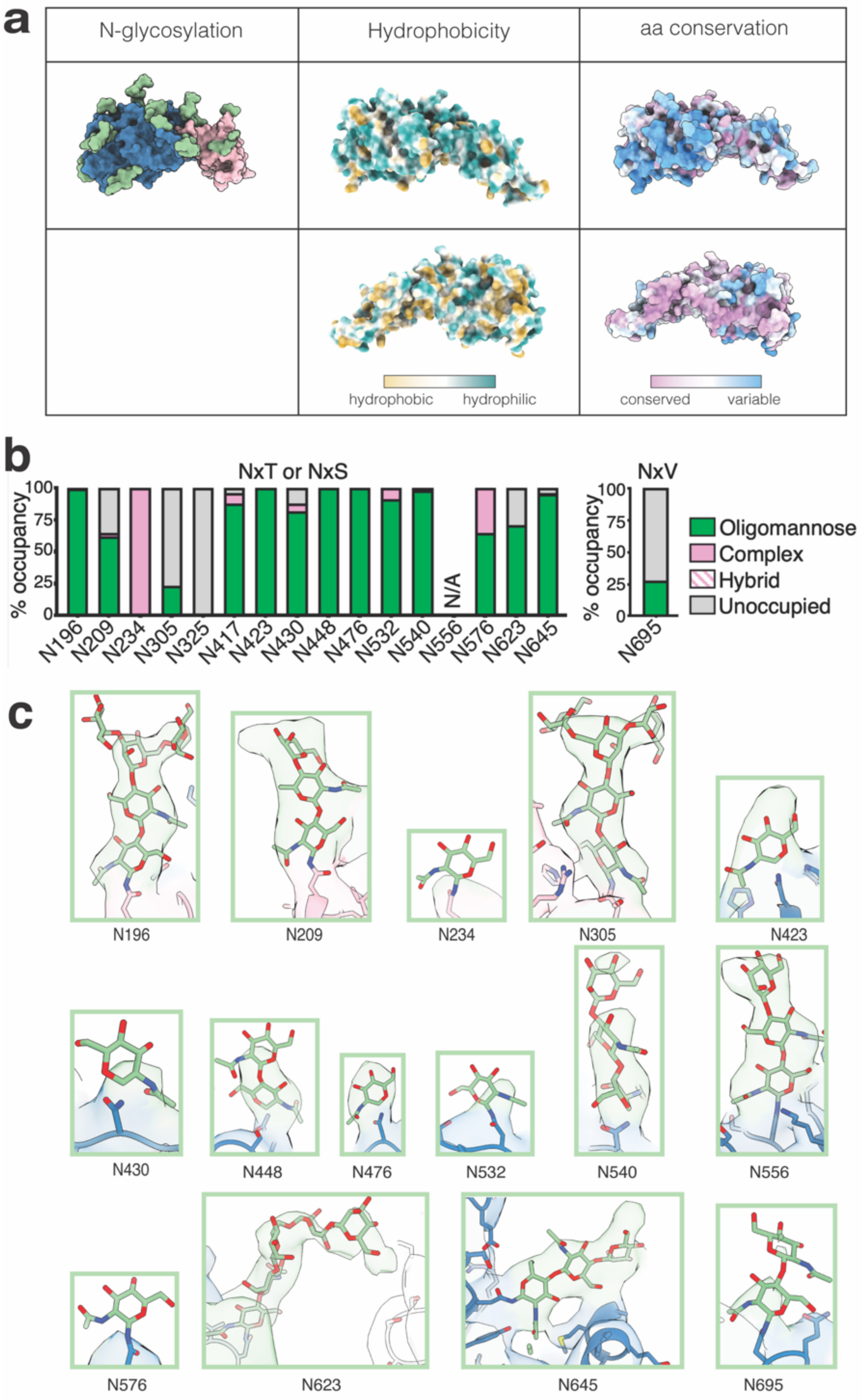
a
N-glycosylation
Hydrophobicity
aa conservation
hydrophobic
hydrophilic
conserved
variable
b
NxT or NxS
NxV
% occupancy
100
75
50
25
0
N196
N209
N234
N305
N325
N417
N423
N430
N448
N476
N532
N540
N556
N576
N623
N645
N/A
N695
Oligomannose
Complex
Hybrid
Unoccupied
c
N196
N209
N234
N305
N423
N430
N448
N476
N532
N540
N556
N576
N623
N645
N695

**Extended Data Fig. 5**. **Glycosylation analysis of E1E2 glycoprotein complex**. (**a**) N-glycosylation sites (in green) on the surface of E1 (pink) and E2 (blue) (left). Hydrophobicity of the surface exposed amino acids in E1 and E2 using the Kyte-Doolittle hydrophobicity scale (middle). Sequence conservation of surface exposed amino acids of E1E2 (right). Top row is 180° rotated compared to the bottom row. (**b**) Relative quantification of distinct glycan types of full-length AMS0232 E1E2 glycoprotein complex determined by LC-MS. The bar graphs show the relative percentage of the glycan processing state at a particular site. Oligomannose-type glycans are shown in green, hybrid in dashed pink, complex glycans in pink and unoccupied sites in gray. N/A, data not available. (**c**) Glycans densities (green) at shown at threshold of 0.19 in ChimeraX. The unsharpened non-focused map of the full complex that produced the best density for E2 stem residues 708-717, as labeled in the PDB deposition, or Best E1E2 Stem map, in Extended Data Fig. 2, was used to generate this visualization. We observed N576 is visible at low signal-to-noise thresholds, while N325 is not. N417 resides in the AS412 region, which remains unresolved in our structure. E1 in shown in pink and E2 is shown in steel blue.

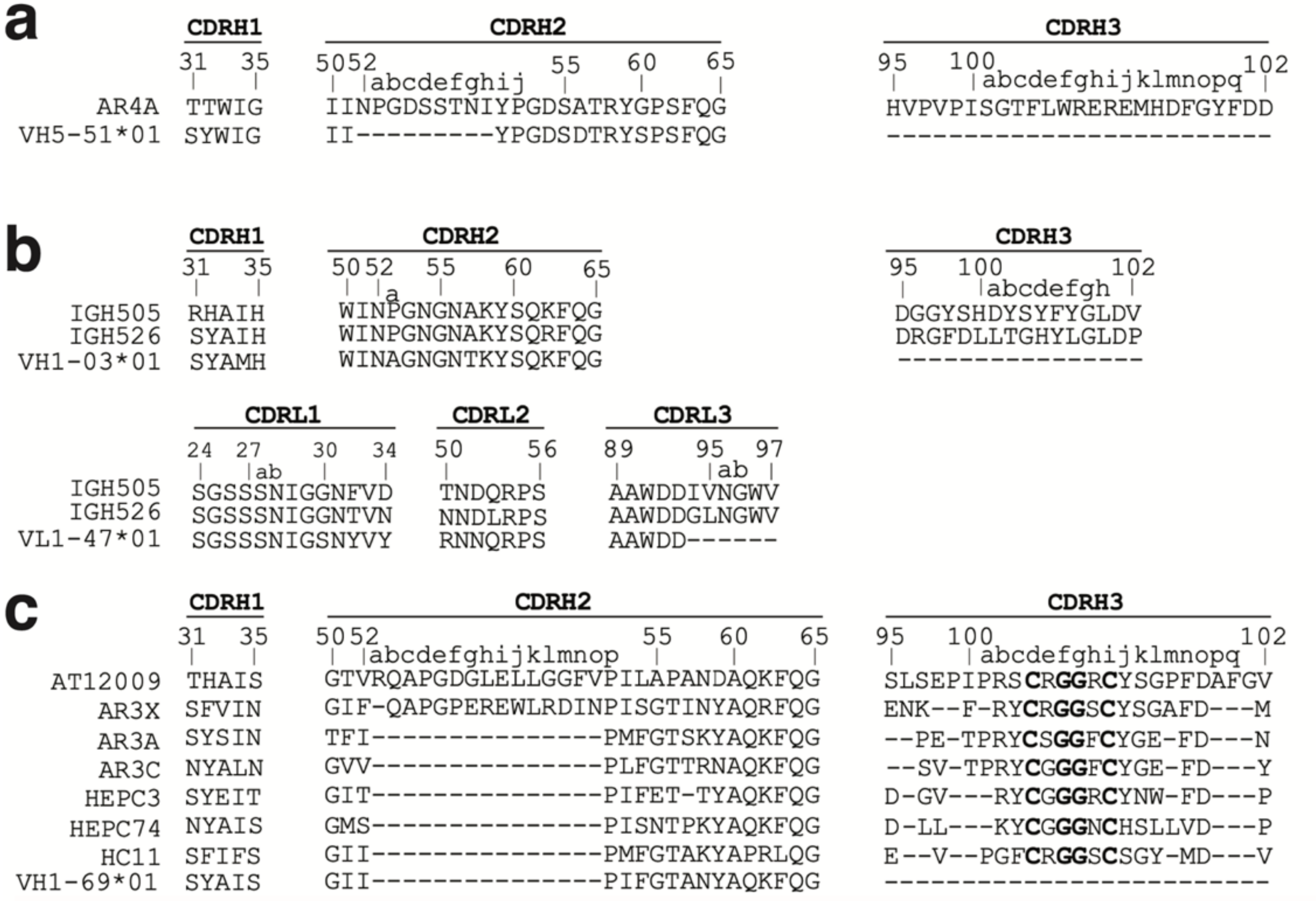

**Extended Data Fig. 6. Alignment of the bNAbs AR4A, AT12009 and IGH505 to their inferred germline.** (**a**). Alignment of the CDRH1-3 of AR4A with its inferred VH5-51*01 germline gene. (**b**). Alignment of the CDRH1-3 and CDRL1-3 of IGH505 and IGH526 with their respective inferred germline genes. (**c**) Alignment of the CDRH1-3 of AT12009 with related AR3-targeting bNAbs that harbor the CxGGxC motif in their CDRH3 and VH1-69*01. CDR annotation and antibody numbering according to Kabat.

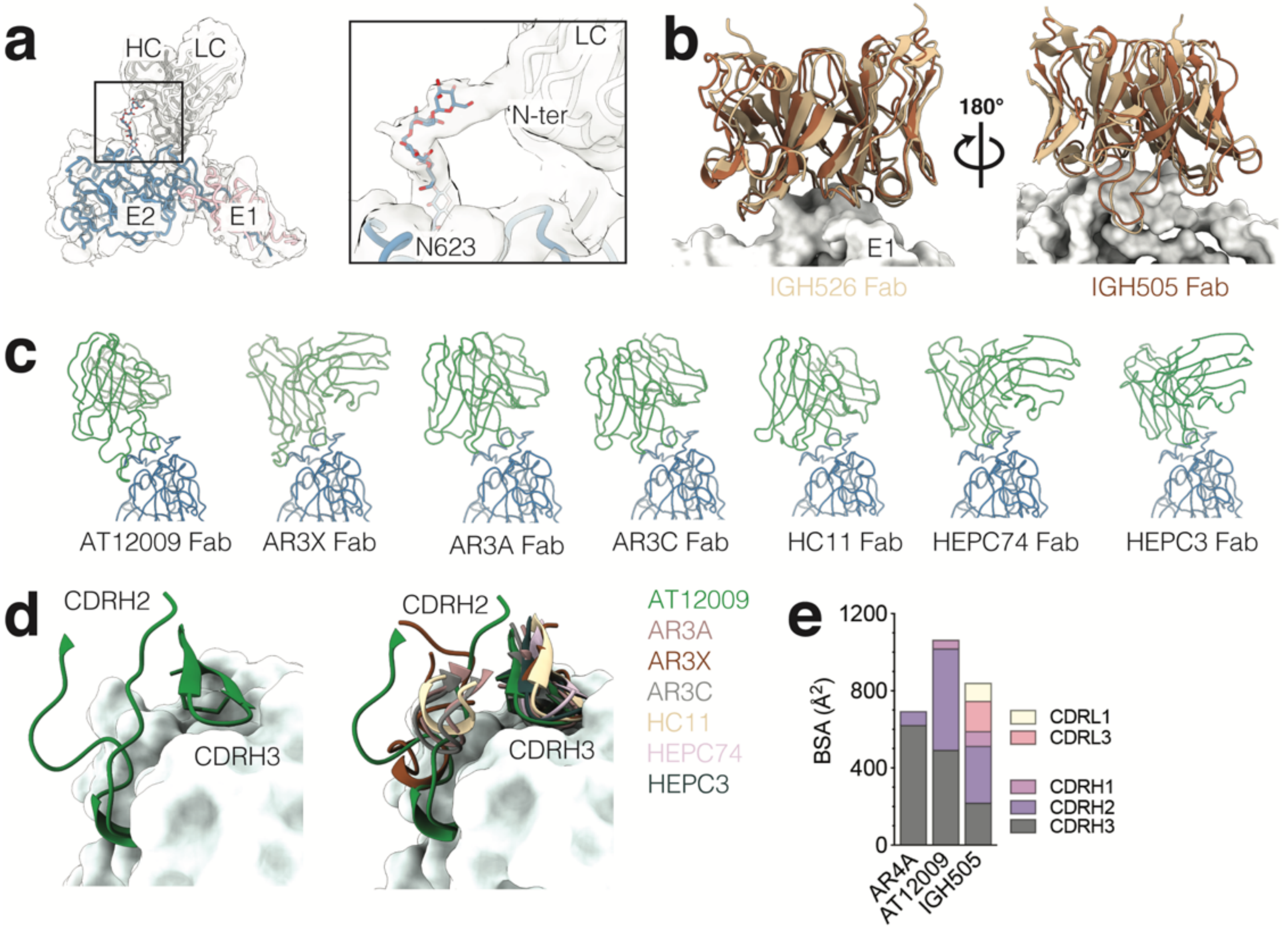

**Extended Data Fig. 7. AT12009 and IGH505 share epitopes with previously known HCV bNAb antibodies.** (**a**) While AT12009 and IGH505 only interact with E2 and E1 peptide region respectively, AR4A Fab not only interacts with the E2 peptide (Fig. 4b) but also with the glycan N623. The unsharpened map is shown in white with the N623 glycan moieties shown in blue and colored by heteroatom interacting with the N-ter domain of the light chain. (**b**) The IGH505 Fab targets the same epitope as IGH526 (PDB 4N0Y) and adopts the same angle of approach. The IGH505 and IGH526 bNAbs are overlayed and shown in brown and yellow respectively. (**c**) CDRH3 motif in E2 front layer-specific HCV bNAbs adopt different orientations. Fab structures in liganded state of AT12009, AR3X (PDB 6URH), AR3A (PDB 6UYM), AR3C (PDB 6UYD), HC11 (PDB 6W04), HEPC74 (PDB 6MEH) and HEPC3 (PDB 6MEJ) are shown. The structures were superimposed on their E2 subunit. Protein backbones are colored green for Fabs and blue for E2 and shown as ribbons. (**d**) The AT12009 ultralong CDRH2 and CDRH3 loops are shown in green laying on the front layer of E2. CDRH2-3 of all HCV bNAb antibodies that target the front layer of E2 are superimposed and shown in different colors. (**e**) Buried surface area between E1E2 and the CDR loops of the AR4A, IGH505 and AT12009 bNAbs.

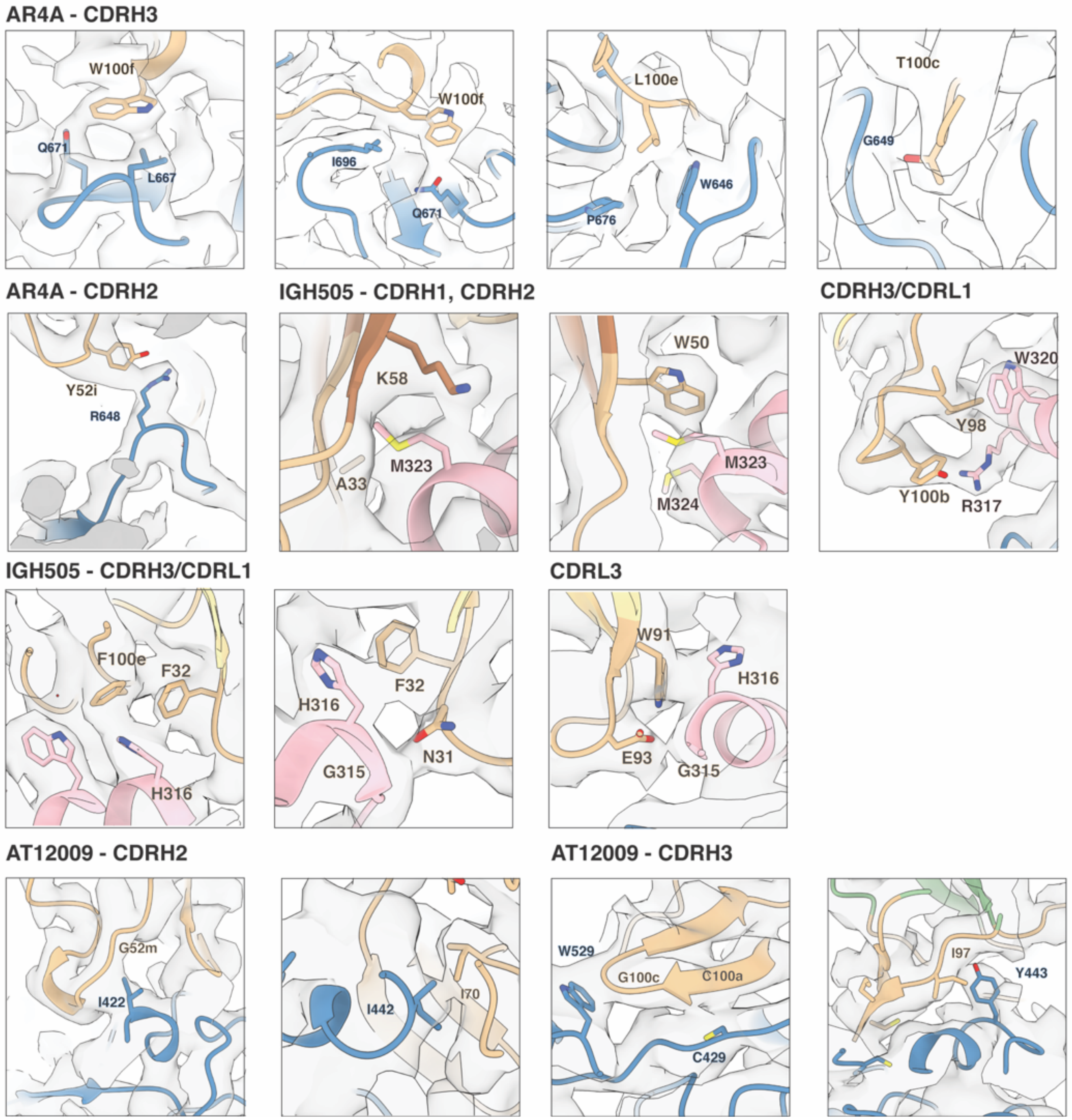

**Extended Data Fig. 8. Close-up views of the amino acid residues involved in the epitope/paratope between E1E2 glycoprotein complex and AR4A, AT12009 and IGH505 Fabs.** Models (ribbon representation) and maps (light gray) are shown and close-up views of each structure with the most relevant epitope/paratope amino acids are indicated. Contact densities are shown at a threshold of 0.1 in ChimeraX.

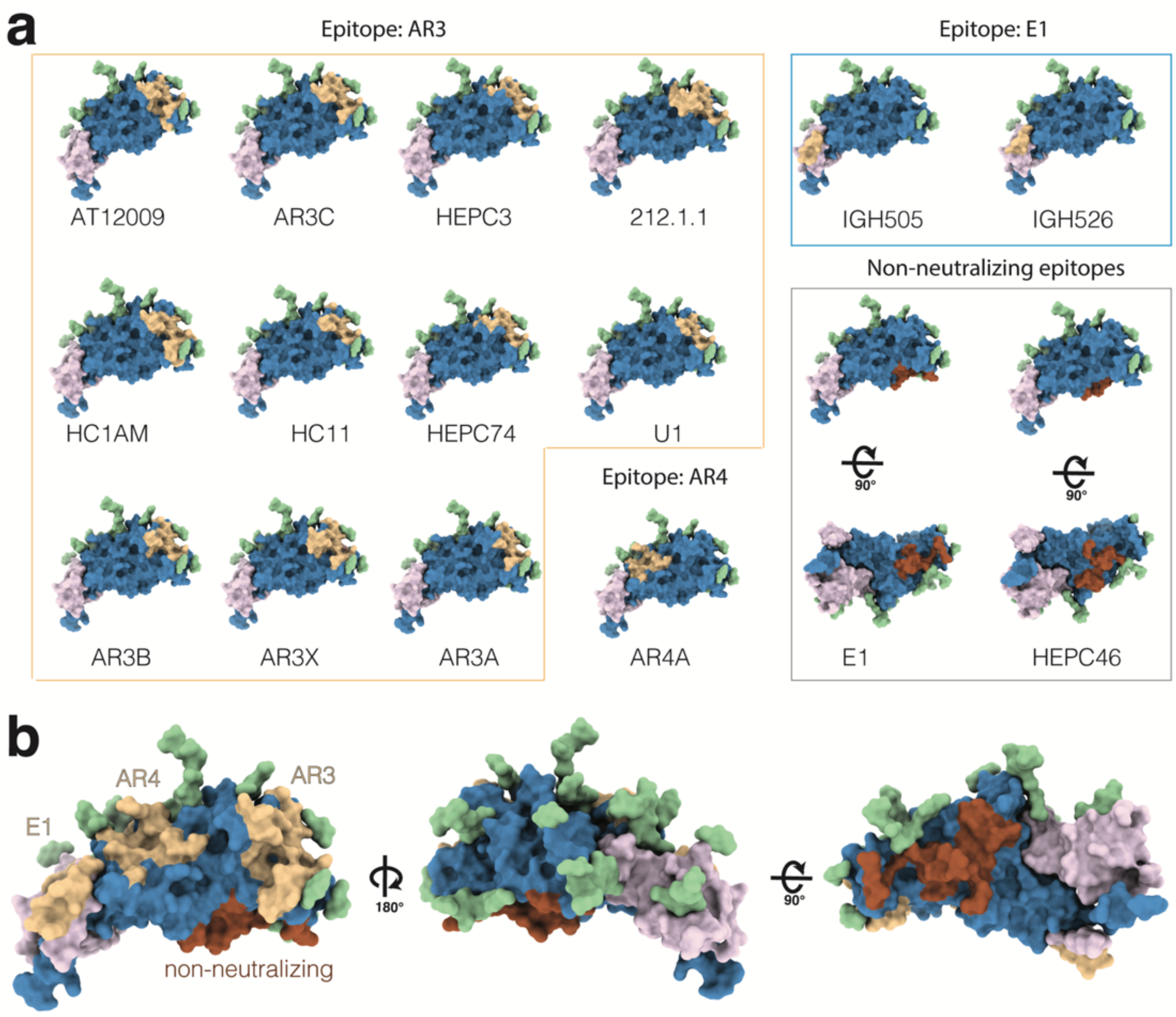

**Extended Data Fig. 9. Footprints of the epitopes of human NAbs and non-NAbs targeting E1E2 glycoprotein complex.** (**a**) Comparison of antigenic region 3 (AR3) and antigenic region 4 (AR4) on E2, a neutralizing epitope on E1 and non-neutralizing epitopes. Footprints were defined on the E1E2 glycoprotein complex as atoms within 4 Å of the indicated Fab. Footprints of neutralizing antibodies are colored in wheat and non-neutralizing antibodies in brown. E2 is colored blue and E1 in pink. (**b**) Summary representation of the footprints of the NAbs and non-NAbs in a E1E2 glycoprotein complex defining a neutralizing face (left), glycan face (middle) and non-neutralizing face (right).
